# A mathematical multi-organ model for bidirectional epithelial-mesenchymal transitions in the metastatic spread of cancer

**DOI:** 10.1101/745547

**Authors:** Linnea C. Franssen, Mark A.J. Chaplain

## Abstract

Cancer invasion and metastatic spread to secondary sites in the body are facilitated by a complex interplay between cancer cells of different phenotypes and their microenvironment. A trade-off between the cancer cells’ ability to invade the tissue and to metastasise, and their ability to proliferate has been observed. This gives rise to the classification of cancer cells into those of *mesenchymal* and *epithelial* phenotype, respectively. Additionally, mixed phenotypic states between these two extremes exist. Cancer cells can transit between these states via *epithelial-mesenchymal transition* (EMT) and the reverse process, *mesenchymal-epithelial transition* (MET). These processes are crucial both for the local tissue invasion and the metastatic spread of cancer cells. To shed light on the role of these phenotypic states and the transitions between them in the invasive and metastatic process, we extend our recently published multi-grid, hybrid, individual-based mathematical metastasis framework (Franssen et al., 2019a). In addition to cancer cells of epithelial and of mesenchymal phenotype, we now also include those of an intermediate *partial-EMT* phenotype. Furthermore, we allow for the switching between these phenotypic states via EMT and MET at the biologically appropriate steps of the invasion-metastasis cascade. We also account for the likelihood of spread of cancer cells to the various secondary sites and differentiate between the tissues of the organs involved in our simulations. Finally, we consider the maladaptation of metastasised cancer cells to the new tumour microenvironment at secondary sites as well as the immune response at these sites by accounting for cancer cell dormancy and death. This way, we create a first mathematical multi-organ model that explicitly accounts for EMT-processes in individual cancer cells in the context of the invasion-metastasis cascade.

## 1. Introduction

To elucidate the process by which a subset of cancer cells from a primary tumour invade the local tissue and spread to distant sites in the body, which is also known as the *invasion-metastasis cascade*, we proposed a first explicitly spatial mathematical modelling framework in Franssen et al. (2019a). The framework described the metastatic process by taking into account the spatiotemporal evolution of individual cancer cells. The motivation for developing such a model is the fact that over 90% of cancer-related deaths arise due to metastatic spread rather than as a consequence of tumour growth at primary sites. Mathematical models can enhance our understanding of the mechanisms underlying biological phenomena. However, with regards to existing models of the invasion-metastasis cascade, we found the following common short-coming: Metastatic spread is an inherently spatial, cell-based physiological process. Yet, previous mathematical models did not capture individual cell dynamics during the invasion-metastasis cascade through a spatially explicit approach, e.g. Iwata et al. (2000); Scott et al. (2013); Cisneros & Newman (2014); Margarit & Romanelli (2016); Iwata et al. (2000); Benzekry et al. (2016); *cf.* literature review in Franssen et al. (2019a). Biologically accurate computational models of the invasion and the secondary metastatic spread of individual cancer cells could be used in a clinical setting to enhance treatment through patient-specific disease predictions (see Figure 1). This is because such models allow to account for e.g. the phenotypic traits of the cells that a tumour consists of in a specific patient, the tumour size and shape, and the tumour microenvironment through the initial conditions of simulations. As explained in Figure 1, the *in silico* simulation results could therefore support clinicians in tailoring treatment to each patient by predicting the individual’s disease evolution, and by testing and optimising treatments in an ethical, time-effective approach.

**Figure 1.**
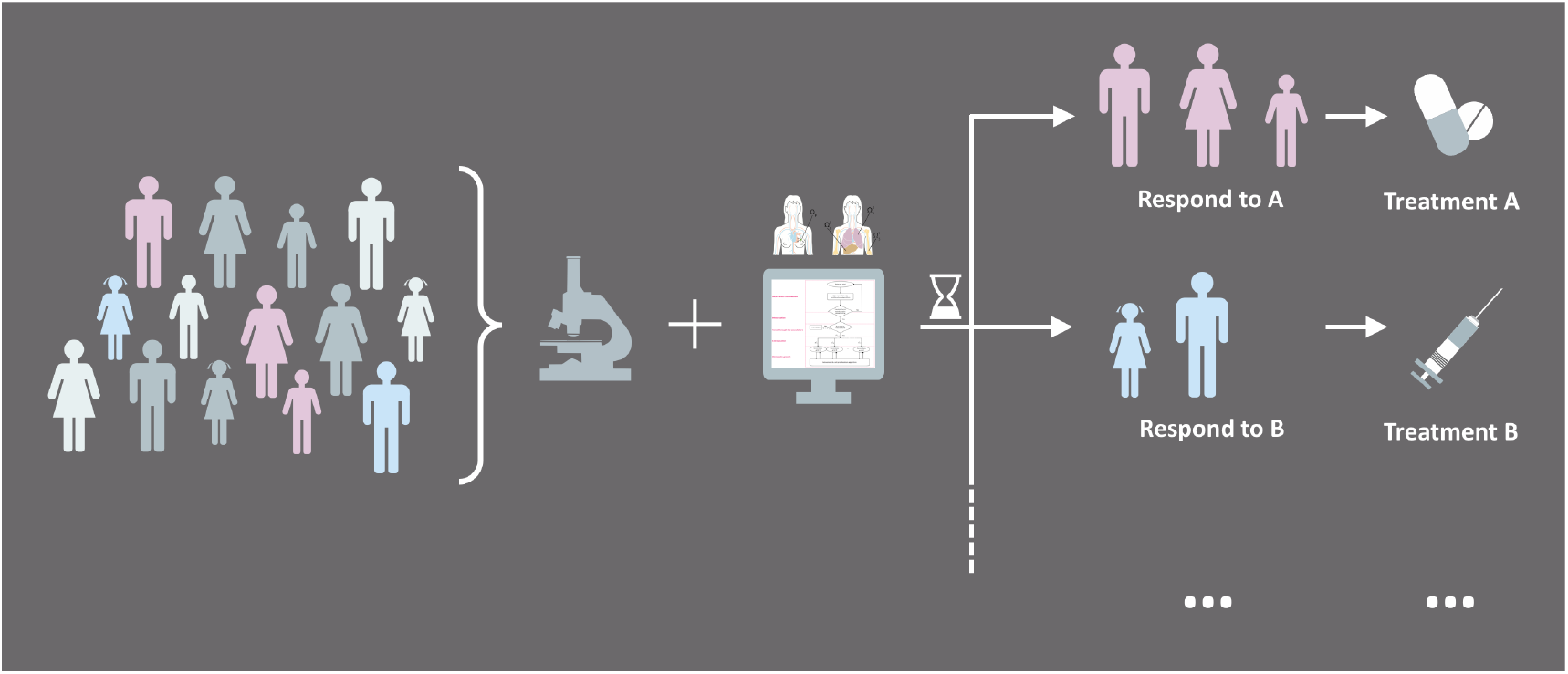
Personalising cancer treatment through computational modelling. In a diverse cohort of cancer patients, different patients respond to different treatment approaches. Modern imaging technologies, methods for image analysis and staining techniques can be combined with spatial, cell-based computational models like the one proposed in this paper. This could, in the future, aid clinicians in identifying the best available treatment for each cancer patient.

To develop a model that will ultimately be used to inform clinicians with regards to cancer treatment decisions, in the concluding section of Franssen et al. (2019a) we proposed several features to enhance the base modelling framework developed therein. *Inter alia*, we concluded that—since *epithelial-mesenchymal transition* (EMT) and *mesenchymal-epithelial transition* (MET) are major factors in the metastatic process—a first natural extension to the modelling framework would be their inclusion. This is achieved in this paper by accounting for phenotypic switching to represent permanent and transient transitions between the epithelial and the mesenchymal phenotypic state as well as a mixed epithelial/mesenchymal state, as biologically observed during the invasion-metastasis cascade (Celià-Terrassa et al., 2012). Thus, in this paper, EMT-related features are introduced to the existing metastasis frame-work to accurately represent their role in the invasion-metastasis cascade. We also account for differences in the *extracellular matrix* (ECM) density of the tissue of the primary and secondary tumour growth sites involved in our simulations. This way, we address another enhancement to the model suggested in Franssen et al. (2019a) by taking a further step towards developing a biologically accurate multi-organ spatially explicit model. Finally, we include the immune response at secondary sites through the modelling of dormancy and death of metastasised cancer cells.

The remainder of the paper is organised as follows. In Section 2, we introduce the biological background of EMT, MET and of the cancer cell phenotypes involved in the context of the invasion-metastasis cascade. In Section 3, we describe how EMT and MET—which we may for simplicity also jointly refer to as the *EMT process* throughout this paper—are included in our general mathematical modelling framework of metastatic spread. As part of this, we give an overview of previous models concerned with EMT in cancer invasion at the beginning of this section. In Section 4, we outline how the computational simulations are set up. In Section 5, we present the simulation results. Finally, in Section 6, we discuss how our results fit in with current biological findings and hypotheses. We also give an overview of future work.

## 2. Biological background

Mutations of key genes in only a few epithelial cells in the body can ultimately lead to the formation of carcinomas, which are the group of solid tumours arising from epithelial tissues in the body. Abnormally rapid proliferation caused by these mutations can result in the formation of an avascular tumour with a diameter of up to approximately 0.1–0.2 cm (Folkman, 1990). Nutrient and oxygen supply to tumour cells in this early avascular stage occurs via diffusion from a vessel source only. The diffusion limit of oxygen is 100–200 µm. Hence, vascular growth is restricted by the metabolic needs of the cells forming the rapidly expanding tumour. It has been observed that, once the tumour growth limit of the avascular phase is reached, a subset of cancer cells start invading the tissue surrounding the tumour either as individual cancer cells or as cancer cell clusters. These invading cells continue to perform random motion but additionally are driven away from the primary tumour mass by gradients in nutrients, oxygen and in the *extracellular matrix* (ECM). Furthermore, the cancer cells secrete chemicals, collectively known as *tumour angiogenic factors* (TAFs), which start recruiting new blood vessels (Folkman & Klagsbrun, 1987)—a process known as *(tumour-induced) angiogenesis*. The resulting newly established vasculature enables the transport of nutrients and oxygen required for further tumour growth. Also, cancer cells may now intravasate into the newly grown blood vessels, travel through the bloodstream and extravasate at distant sites in the body where space and nutrients are less of a limiting factor to growth. The successful relocation of cancer cells from a primary location to a secondary location in the body via the described sequence of steps in the invasion-metastasis cascade is known as *metastatic spread*. Successfully extravasated cancer cells occur either as single *disseminated tumour cells* (DTCs) or as small clusters of cancer cells, called *micrometastases*. The majority of micrometastases and, even more so, of DTCs die due to maladaptation to the new microenvironment as well as due to the local immune response (Aceto et al., 2014). However, surviving cancer cells may proliferate, which can lead to the formation of secondary tumours, called *metastases*, at sites in the body away from the primary tumour. Other DTCs and micrometastases may remain dormant at first but have the potential to proliferate into vascularised metastases at the secondary sites at some later point in time. The full process we have described here is shown schematically in Figure 2. Its biological background is further explained in Franssen et al. (2019a) by considering each of the steps in the invasion-metastasis cascade—i.e. cancer cell invasion, intravasation, vascular travel, extravasation and regrowth at new sites in the body—in turn.

**Figure 2.**
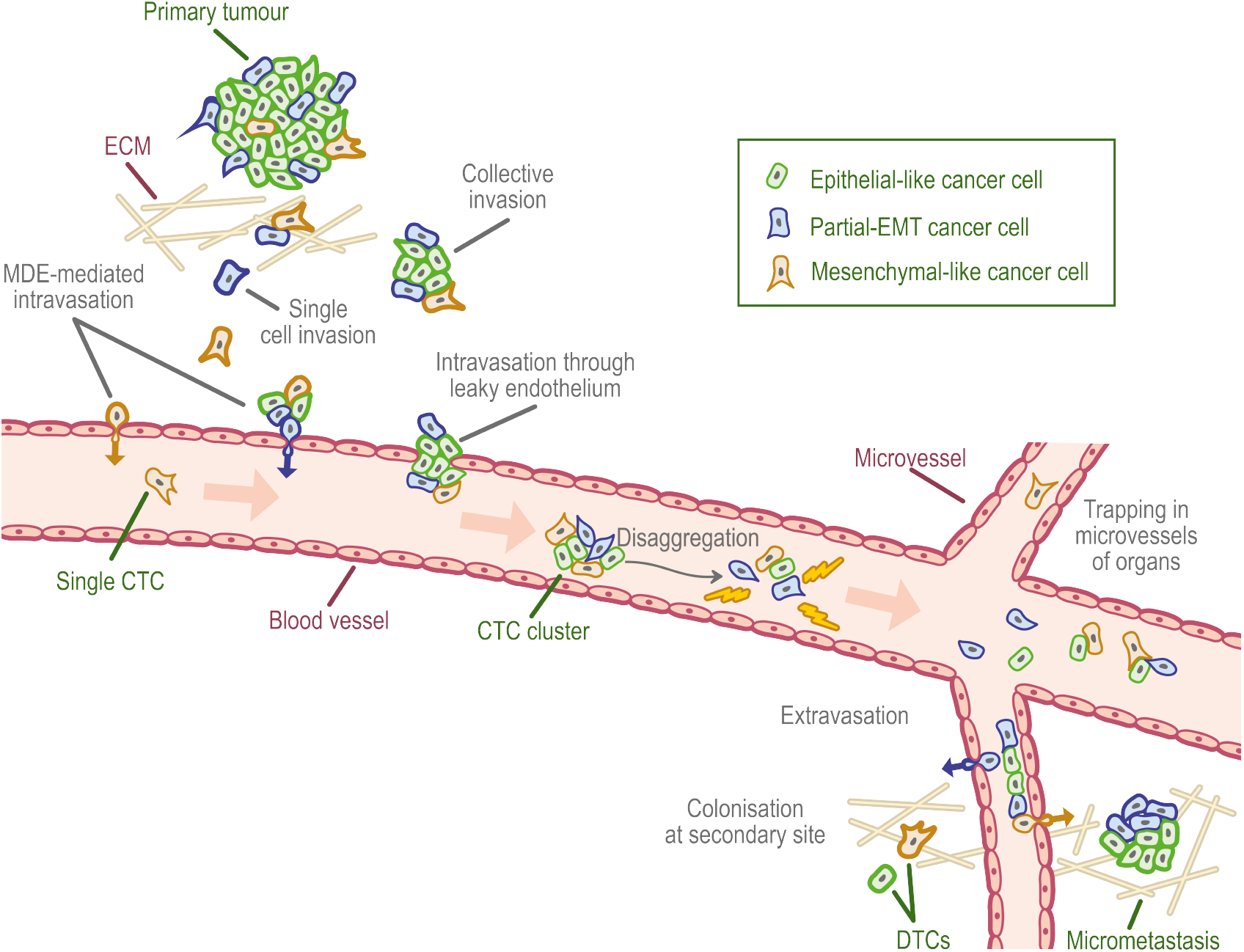
Schematic overview of the invasion-metastasis cascade. Single mesenchymal-like and partial-EMT cancer cells as well as heterogeneous clusters consisting of mesenchymal-like, partial-EMT and epithelial-like cancer cells break free from the primary tumour and invade the surrounding tissue (top left). They can intravasate via active *matrix-degrading enzyme* (MDE)-mediated and passive mechanisms (mid-left, along epithelium of the vessel). Once in the vasculature, *circulating tumour cell* (CTC) clusters may disaggregate (centre) and CTCs may die. Surviving cells may extravasate through the walls of the microvasculature to various secondary sites in the body (bottom right). Successful colonisation there can result in either *disseminated tumour cells* (DTCs) or in micrometastases, which have the potential to develop into full-blown metastases.

Cancer cells adapt to the environmental requirements of the various steps of the invasion-metastasis cascade via changes in phenotype (Jolly et al., 2017a). EMT and MET are a canonical group of—at least transiently—observed phenotypic changes that are assumed to be crucial for metastatic spread (Guo et al., 2012; Ye et al., 2015; Krebs et al., 2017). Various combinations of so-called *EMT-inducing transcription factors* (EMT-TFs) together with a number of extracellular molecules in the tumour microenvironment and related pathways are thought to trigger EMT (Jie et al., 2017). The cell-cell adhesion between formerly epithelial-like cancer cells is typically reduced upon activation of EMT. At the same time, the cancer cells tend to express more cell-matrix adhesion enhancing molecules like cadherin (Micalizzi et al., 2010). As part of this combination of changes, the characteristic polygonal cobblestone-like cell shape of epithelial cells is progressively replaced by a spindle-shaped morphology, as shown on the right of Figure 3. Also, the motility and invasiveness of the cancer cells are enhanced (Jie et al., 2017; Dongre & Weinberg, 2019). As another result of EMT, the cells become increasingly potent at degrading the underlying basement membranes of organs and vessels as well as the ECM via the expression of *metalloproteases* (MMPs) (Dongre & Weinberg, 2019). As a trade-off, they become less proliferative. MET, can reverse the phenotypic changes induced by EMT, thus—generally speaking—causing the cells to become less motile and invasive while enhancing their proliferative potential.

**Figure 3.**
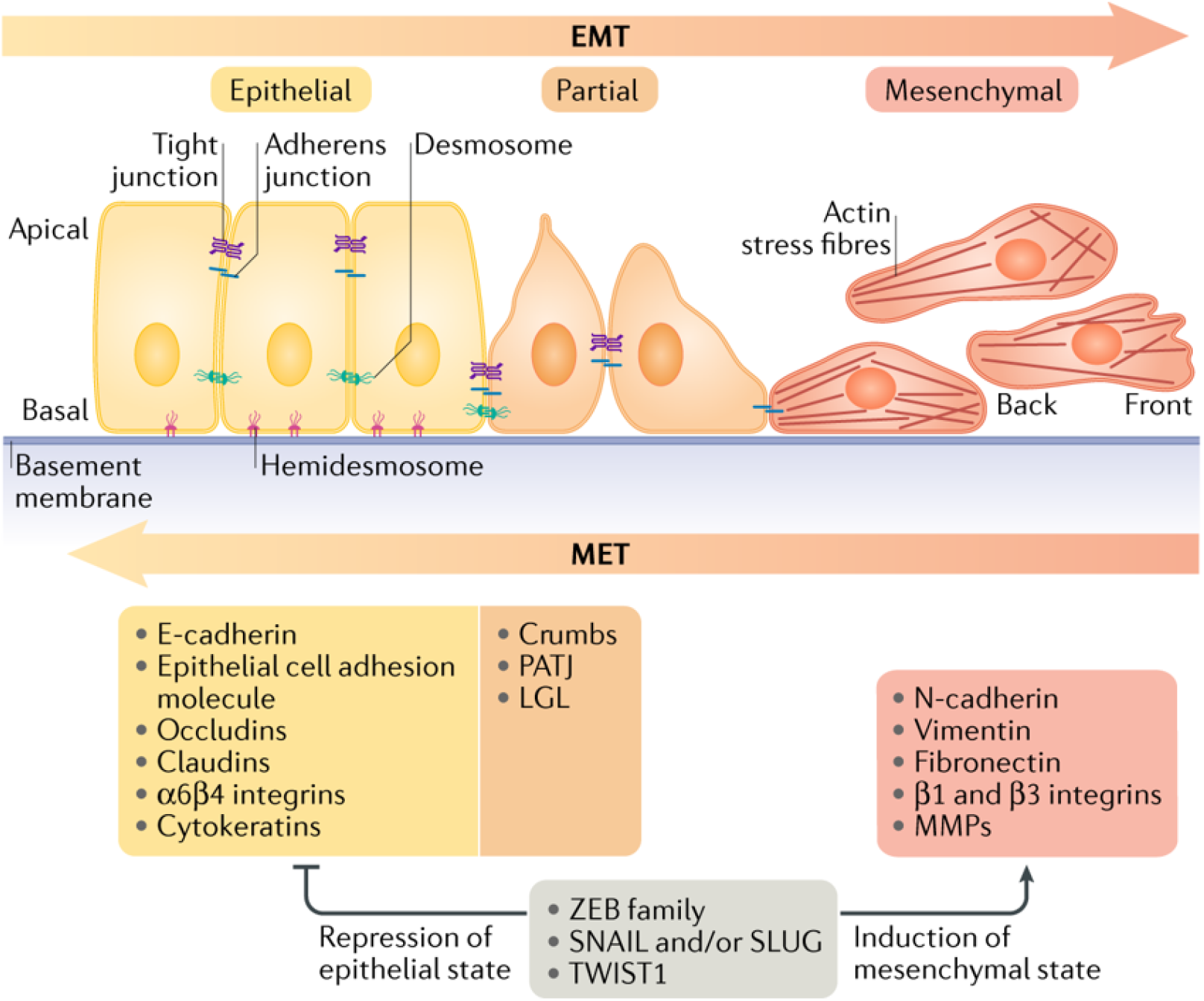
Typical EMT transition states in carcinoma progression and metastatic spread. During carcinoma invasion and metastatic spread, the formerly epithelial phenotype of cancer cells (left) can—via the processes of partial or full EMT—evolve to a partial-EMT (middle) or a mesenchymal (right) phenotype. Generally, the further to the right of the EMT spectrum a cell is located, the less it attaches to other cells. The trade-off for this gain in motility is a decreased proliferative potential. MET is the reverse process. It is indicated by the arrow along the bottom of the figure. Reproduced from Dongre & Weinberg (2019) with permission from Nature Publishing Group.

Traditionally, the EMT-process has been viewed to result in cells of epithelial and of mesenchymal phenotype in a binary sense (Pastushenko & Blanpain, 2018; Dongre & Weinberg, 2019). Yet, more recently, intermediate states—commonly referred to as *hybrid*, *incomplete* or *partial-EMT* states—on the spectrum between the fully epithelial and fully mesenchymal state have been shown to exists in various cell lines of patient xenografts and of human primary cancers, such as breast, head and neck, and pancreatic cancer (Pastushenko & Blanpain, 2018). Cancer cells in these intermediate phenotypic states are assumed to show a variety of combinations of the above-mentioned phenotypic traits. The full transition from an epithelial to a mesenchymal state, which had formerly been assumed to be the only possible outcome of EMT, has recently been shown to actually be rare during carcinogenesis (Dongre & Weinberg, 2019). Furthermore, cell cycle arrest may occur in fully mesenchymal cancer cells (Vega et al., 2004; Lovisa et al., 2015), while partial-EMT cancer cells continue to be able to proliferate (Handler et al., 2018).

In what follows, the roles of EMT and MET as well as of epithelial-like, partial-EMT and mesenchymal-like cancer cells during the various steps of the invasion-metastasis cascade are elucidated in more detail. The five steps of the invasion-metastasis cascade are printed in bold for better orientation. A more in-depth description of the EMT-unrelated features of these steps of the invasion-metastasis cascade is provided in Franssen et al. (2019a).

### Local cancer invasion

Carcinomas are tumours that arise from epithelial tissue. However, cancer cells have been found to either invade as single cells of partial-EMT or of mesenchymal phenotype or as clusters, which often consist of cancer cells of heterogeneous phenotypes (Friedl & Wolf, 2003). Hence, EMT of some degree—at least in a subset of the cancer cells—at the primary site is a prerequisite for this first step of the invasion-metastasis cascade (Francart et al., 2018; Pastushenko & Blanpain, 2018). Migrating cells usually employ their acquired mesenchymal traits, i.e. the decrease or loss in cell-cell adhesion and increases in cell-ECM adhesion and in matrix-degrading enzyme (MDE)-expression, to invade (Friedl & Wolf, 2003; Bill & Christofori, 2015). This hypothesis is, for example, supported by reports suggesting that invading cancer cell clusters contain cells that have undergone partial EMT *in vivo* (Tsai et al., 2012; Ocaña et al., 2012). Moreover, the occurrence of *clusters* highlights that partial EMT allows for the cancer cells to maintain at least some aspects of epithelial cell-cell adhesion (Cheung & Ewald, 2016). Furthermore, the spatial location of cancer cells of partial-EMT and of epithelial phenotype was investigated by Puram et al. (2017) *in situ* in oral cavity *head and neck squamous cell carcinomas* (HNSCCs). Using immunohistochemistry to stain a collection of tumours, they found that, while the core of the tumours contained malignant cells of epithelial phenotype, partial EMT had occurred in the cancer cells at the leading tumour edge in the proximity of *cancer-associated fibroblasts* (CAFs) in the tumour microenvironment. A corresponding explanation in the form of a diagram and a stained tissue sample is shown in Figure 4.

**Figure 4.**
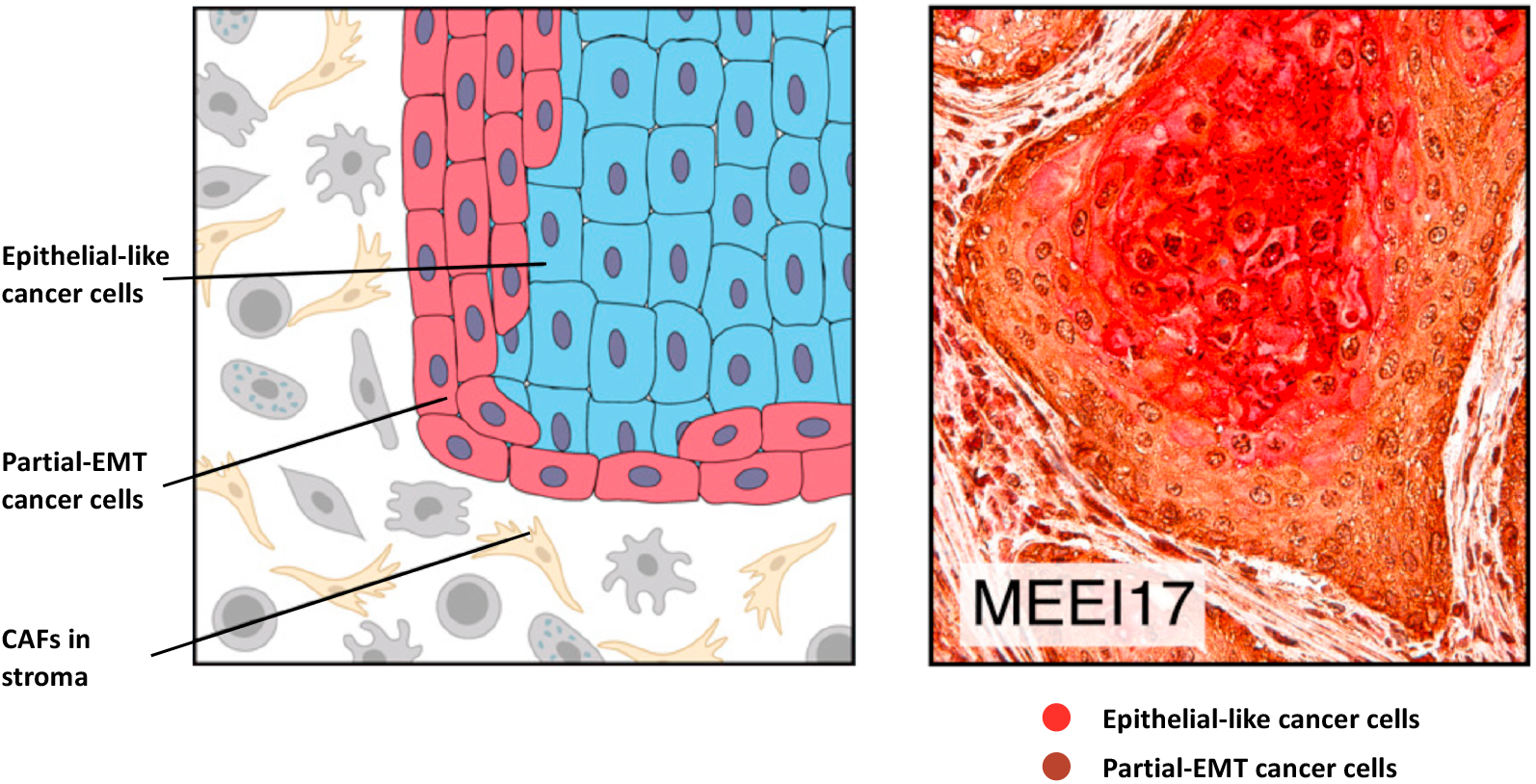
Partial EMT occurs at leading tumour edge in HNSCC. In situ spatial location of cancer cells expressing a partial EMT programme versus those of epithelial phenotype within HNSCC tumours, both schematically (left) and in human tissue (right). On the left, immunohistochemistry was used to stain the tumour for PDPN, one of the top genes in the partial-EMT programme, as well as for SPRR1B, an epithelial differentiation marker. Partial-EMT cancer cells were located at the leading edge of tumours in proximity to *cancer-associated fibroblasts* (CAFs) in the surrounding stroma, epithelial-like cancer cells at the core of tumours. Reproduced from Puram et al. (2017) with permission from Elsevier.

### Intravasation

As explained in detail in Franssen et al. (2019a), unless vessels are ruptured—for instance as a consequence of the tumour microenvironment modulating effects of EMT-TFs (Jolly et al., 2017b)—and subsequently opportunistically entered by cancer cells, MDE-expressing cancer cells are required to allow the intravasation of cancer cells into the vasculature. Therefore, epithelial-like cancer cells cannot gain access to undamaged vessels, while partial-EMT and mesenchymal-like cancer cells can (Jolly et al., 2018). Similarly, cancer cell clusters that consist at least partially of partial-EMT or mesenchymal-like cancer cells can enter undamaged vessels using MDEs.

### Travel through the vasculature

The majority of *circulating tumour cells* (CTCs) and CTC clusters in the vasculature were found to be of partial-EMT phenotype (Jolly et al., 2018). Armstrong et al. (2011) found that in women with metastatic breast cancer and men with castration-resistant prostate cancer more than 75% and 80% of CTCs, respectively, co-expressed epithelial and mesenchymal markers. Similarly, studies by Thiery & Lim (2013) and by Yu et al. (2013) reported that a significant proportion of CTCs was of partial-EMT or mesenchymal-like phenotype in patients with metastatic breast cancer. CTCs of partial-EMT phenotypes have further been observed in the blood of patients with cancer of the liver, prostate and lungs as well as in patients with colorectal, nasopharyngeal and gastric cancer. In these types of cancer, the partial-EMT phenotype correlates with poor clinical prognosis when compared to the occurrence of cancer cells of purely epithelial or purely mesenchymal phenotype (Pastushenko & Blanpain, 2018). The prominence of partial-EMT cancer cells at the tumour edge as well as their ability to intravasate into the vasculature using MDEs offer potential explanations for these findings. An additional explanation is that at least a subset of partial-EMT CTCs is more resistant to anoikis, i.e. to apoptosis induced by lack of correct cell-ECM attachment (Huang et al., 2013). However, independent of phenotype, both single CTCs and CTC clusters are exposed to physical stresses and to attacks by natural killer cells in the vasculature. One consequence is that only a small number of cancer cells shed from the primary tumour actually reach the microvasculature at metastatic sites. Another effect of this is the (partial) disaggregation of cancer cell clusters, as shown in the centre of Figure 2. This leads to smaller CTC clusters and to an increased number of single CTCs.

### Extravasation

For the subset of CTCs and CTC clusters that do reach the microvessels at distant organs, extravasation, i.e. the translocation from the vasculature to the tissue at a secondary site, is the next step. During extravasation, the phenotype of the cancer cells is believed to play a tangential role at most—CTCs of all phenotypes appear to be able to extravasate (Banyard & Bielenberg, 2015) with the aid of mechanisms explained further in Lambert et al. (2017); Franssen et al. (2019a).

### Colonisation and metastatic growth

However, cancer cell phenotypes are, once again, of crucial importance when it comes to the colonisation and metastatic growth of cancer cells at the secondary sites. Also, EMT alone fails to explain this last step of the invasion-metastasis cascade, given that macrometastases in humans often present similar histopathological traits to the primary tumours they originate from. These traits include a mainly epithelial-like morphology (Pastushenko & Blanpain, 2018) with a relatively small subset of cancer cells with phenotypes further along the EMT spectrum (Dongre & Weinberg, 2019)—despite the above-described evidence of the abundance of partial-EMT CTCs in the vasculature. Consequently, this suggests that some degree of MET is needed for macrometastatic growth. A murine prostate cancer model by Ruscetti et al. (2015) delivers insight into this. Cancer cells in macrometastases that had spread to the lungs were found to have mainly epithelial markers and few mesenchymal markers; the inverse constitution was found in dormant micrometastatic lesions. Coherently, in a study by Ocaña et al. (2012), it was proposed that the constant overexpression of the EMT-inducer PRRX1 in human breast tumour cell lines, which were injected intravenously into chick embryos, may lock cancer cells in a mesenchymal-like phenotypic state. This was suggested to inhibit the cells from performing MET, which, in turn, failed to give rise to lung metastases. Similarly, Kröger et al. (2019) concluded from several studies that a stable mesenchymal-like phenotype without any MET potential cannot succeed in metastatic re-seeding.

Finally, as elaborated in Franssen et al. (2019a), it is noteworthy that experimental evidence suggests that less than 0.07% of all initially intravasated single CTCs form micrometastases and less than 0.018% ± 0.017% form macrometastases 13 days after intravasation (Luzzi et al., 1998; Valastyan & Weinberg, 2011). Maladaptation to the new tumour microenvironment, with the consequence that only a few cells are able to proliferate, is regarded to be a main contributor to the poor survival at secondary sites (Dongre & Weinberg, 2019). Compared to single CTCs, CTC clusters were described to have 23 to 50 times the metastatic potential (Aceto et al., 2014). One explaining factor for this is that cell heterogeneity, as often found in such clusters, can be advantageous during colonisation (Jolly et al., 2018).

## 3. The EMT/MET multi-organ extension of the metastasis modelling framework

In this section, the inclusion of EMT-related processes into the recently introduced mathematical frame-work for the modelling of the metastatic spread of cancer by Franssen et al. (2019a) is outlined. Only new EMT-related features will be established here—the reader is referred to Appendix A to consult the existing underlying metastasis modelling framework, onto which we impose the alterations described in this section. Further, we introduce changes to the existing framework that allow us to differentiate between the cell behaviour on the various organs as well as to account for dormancy and death of metastasised cancer cells as a result of the potential immune response at and maladaptation to secondary sites. We begin by giving an overview of existing mathematical models that include, in the wider sense, EMT-related features in the context of spatially explicit cancer invasion. A review of such mathematical models of the EMT-process in the context of metastasis will be omitted as, to our knowledge, this paper is the first metastasis model to include the roles of EMT and MET, and of the corresponding phenotypes of individual cancer cells in a spatially explicit manner.

Andasari et al. (2011) extended and analysed a system of equations initially proposed in Chaplain & Lolas (2005) to represent the interaction between cancer cells, the MDE *urokinase-type plasminogen activator* (uPA), uPA inhibitors of type PAI-1, the ECM-cleaving and MMP-activating enzyme plasmin, and the ECM component vitronectin. They allowed for cancer cells to mutate into a phenotype which diffuses, migrates and proliferates more rapidly, which was modelled using a Heaviside function. While the current biological evidence on EMT-related phenotypic changes somewhat contradicts the notion of such a ‘go-*and*-grow’ mutation, the proposed model was an important step towards modelling mutations in cell phenotype in spatial cancer invasion models. Gerisch & Chaplain (2008) modified the local haptotaxis-based continuum *partial differential equation* (PDE) model proposed in Anderson et al. (2000) to include cell proliferation and ECM remodelling as well as cell-matrix and cell-cell adhesion. This was achieved using an integro-differential PDE model, which incorporated cell-cell adhesion using integral terms. Domschke et al. (2014) extended this model further. In particular, they introduced a subpopulation of cancer cells that arose from the initial cell population by mutation, again by using a Heaviside function. The mutation resulted in a decrease in self-adhesion of the cancer cells and an increase in cell-matrix adhesion, which caused the mutated cancer cells to spread more rapidly into the surrounding tissue. This is coherent with the current biological understanding that EMT causes more invasive phenotypes. In order to include physiological mechanisms that lead to EMT, Hellmann et al. (2016) modelled EMT from differentiated cancer cells to *cancer stem cells* (CSCs), which have biological properties comparable to the epithelial-like and the mesenchymal-like cancer cells in our model, respectively. In this approach, EMT was triggered by *epidermal growth factors* (EGFs) in the ECM. Subsequently, an advection-reaction-diffusion system of Keller-Segel taxis type was used to study the invasion of both types of cancer cells into the ECM. Numerical simulations were proposed as a proof of concept to show that combining the two systems can account for EMT in a biologically accurate manner. Sfakianakis et al. (2017) developed this model of EGF-driven EMT further. In the corresponding simulations, the detachment of CSCs from the main tumour body of differentiated cancer cells—due to their ability to invade the tissue comparatively more rapidly—was reproduced qualitatively. More recently, Sfakianakis et al. (2018) introduced a coupled two-dimensional hybrid system that governed the spatiotemporal evolution of individual mesenchymal cancer cells by a system of *stochastic differential equations* (SDEs), while the collectively moving epithelial cancer cells, the ECM and the MMPs evolved according to PDEs. This novel modelling technique considered the effects of EMT and MET on cancer invasion using phase transition operators. As a result, the *in silico* invasion assays simulated using the Sfakianakis et al. (2018) approach presented ‘islands’ of invading cancer cells ahead of the expanding initial main cancer cell mass, which had arisen from EMT and subsequent MET. These ‘islands’ away from the tumour mass are frequently observed *in vivo* but do not typically present themselves in solely macroscopic or atomistic cancer invasion models. This model was extended to a three-dimensional setting in Franssen et al. (2019b). Moreover, the resulting model was parametrised to accurately represent organotypic invasion assays of oral squamous cancer cells in an experimental invasion model proposed by Nurmenniemi et al. (2009). Simulations were found to qualitatively and quantitatively agree with the experimental invasion assays.

From the literature review, we draw several conclusions. Firstly, no spatially explicit model that describes the role of EMT and MET in metastatic spread—as opposed to their role in invasion alone— exists. Consequently, none of the existing models capture the site- and location-dependent occurrence of EMT and MET in all of the steps of the invasion-metastasis cascade—i.e. in cancer cell invasion, intravasation, vascular travel, extravasation and during regrowth at new sites in the body—in a spatial manner. Secondly, to our knowledge, the simulations from existing spatiotemporal ECM invasion models that account both for epithelial from mesenchymal cancer cell populations as well as for the transition between the phenotypic states, such as Domschke et al. (2017), lack the inclusion of intermediate partial-EMT phenotypes. Yet, it has recently become evident that cancer cells of partial-EMT phenotype are crucial to the EMT process, as explained in Section 2. With the aim of closing the current gap in the literature, we propose an extension to the spatially explicit hybrid modelling framework in Franssen et al. (2019a). The resulting model describes the invasive growth dynamics both of the primary tumour by—*inter alia*—accounting for EMT, as well as growth in the early avascular stages at potential secondary metastatic sites by accounting for MET. Additionally, transport from primary to secondary sites is modelled. In what follows, we introduce the ideas and assumptions that the EMT extension of the metastasis framework builds on. As Franssen et al. (2019a), we use *G* + 1 non-overlapping spatial domains to represent the primary tumour site, *Ω*_P_ ⊂ ℝ^2^, as well as the *G* ∈ ℕ spatial domains representing the sites of potential secondary metastatic spread, 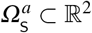, where *a* = 1, 2,…, *G*. As previously, we represent the MMP-2 concentration and the ECM density at position 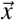 at time *t* in these spatial domains by the continuous functions 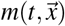 and 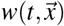, respectively, while capturing the spatiotemporal evolution of epithelial, partial-EMT and mesenchymal cancer cells as well as of the membrane-bound MT1-MMP in a discrete approach, *cf.* Anderson & Chaplain (1998); Anderson et al. (2000); Franssen et al. (2019a). Also analogously to Franssen et al. (2019a), we allow cancer cells to travel from primary to secondary sites via the vasculature by designating locations in the primary spatial domain to function as entry points into blood vessels and, similarly, impose a spatial map of exit locations from the vasculature onto the secondary metastatic domains.

The EMT-related features that are novel to the metastatic framework are explained according to which key step of the invasion-metastasis cascade—i.e. cancer cell invasion, intravasation, vascular travel, extravasation and metastatic growth—they belong to. To enhance the clarity of presentation, as Franssen et al. (2019a), we begin each paragraph by printing the description of corresponding the step in bold. We also label the respective sections in the flowchart presented in Figure 5, which visually describes the model, accordingly.

**Figure 5.**
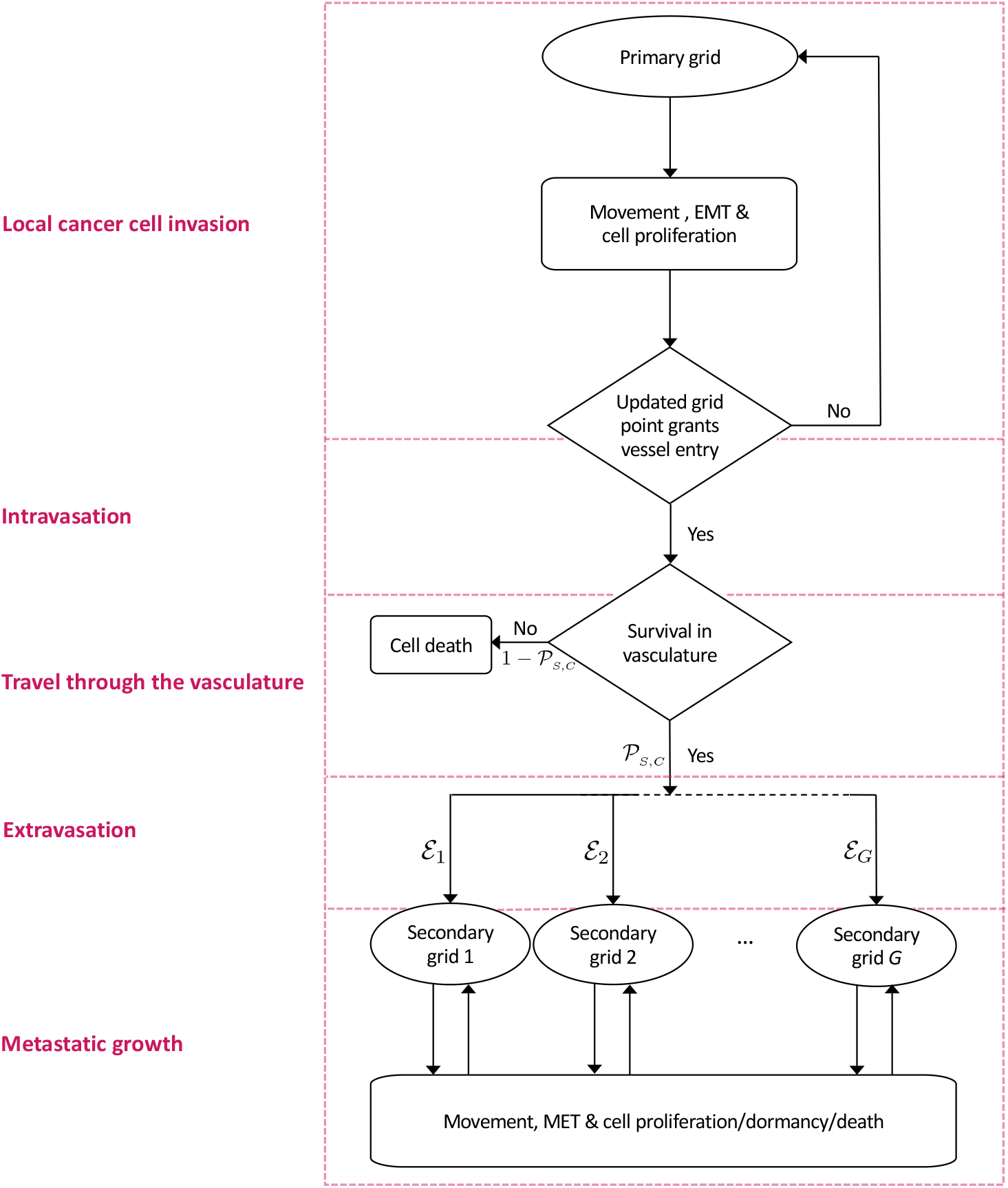
Flowchart of the extended invasion-metastasis hybrid model. At each time step, each cancer cell on the primary grid may move, may perform EMT with some (location-dependent) probability and may proliferate as explained in detail in the text. A cancer cell remains on the primary grid during the respective time step, unless it is placed on a grid point that represents a blood vessel. In the latter case, single CTCs and CTC clusters may enter the vasculature. They spend a number of time steps in the circulation and survive with a probability of 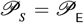, 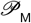 or 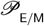 in the case of single CTCs of epithelial, mesenchymal and partial-EMT phenotype, respectively, and with a probability of 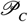 in the case of CTC clusters. Cancer cells that do not survive are removed from the simulation. Surviving CTCs and CTC clusters are placed onto one of *G* secondary grids with the respective probability ℰ_1_, ℰ_2_,…, ℰ_*G*_. Cancer cells on the secondary grids move and proliferate like cancer cells on the primary grid (potentially with different parameter values to represent organ- and patient-specific differences in the local tumour microenvironment). However, partial-EMT cells may now revert to cells of an epithelial phenotype via MET and there furthermore exists a probability for cell death and dormancy. For better orientation, the red boxes with their labels on the left correspond to the sections indicated in bold in Sections 2 and 3 of the text as well as in Appendix A.

**Figure 6.**
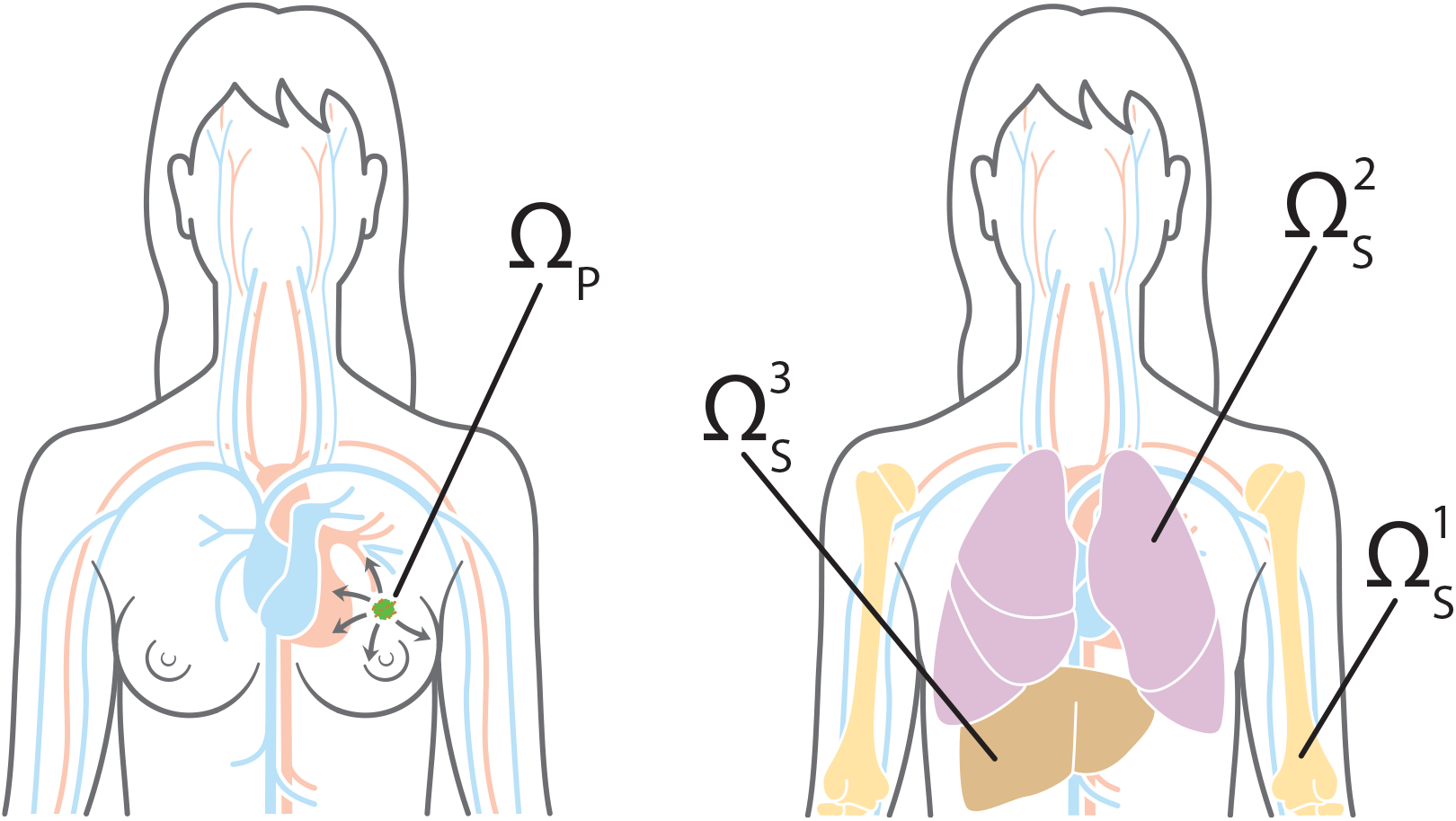
Primary and metastatic sites. To give an example of how the general modelling framework can be applied to a specific clinical setting, we chose the primary site *Ω*_P_ in our simulations to represent the breast (left). Potential secondary metastatic sites 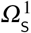, 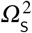, 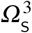 were chosen to represent the bones, the lungs and the liver, respectively (right). Cancer cells can reach the secondary sites by travelling through the blood system.

### Local cancer cell invasion

As explained in detail in Franssen et al. (2019a), the evolution of the MMP-2 concentration and of the ECM density are modelled in a continuum approach. To account for the inclusion of partial-EMT cancer cells in our model, we extend equations (A.1) and (A.2) from the former model slightly. Accordingly, the spatiotemporal evolution of the MMP-2 concentration 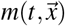 is given by

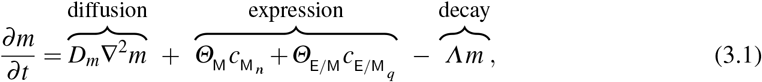

along with zero-flux boundary conditions. Here, 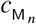, *n* = 0, 1, 2,…, *Q*, and 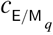, *n* = 0, 1, 2,…, *Q*, with *n* + *q* ≤ *Q*, denote the presence of up to a total of *Q* mesenchymal-like or partial-EMT cancer cells at a given position 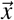, following the notation by Stéphanou et al. (2006) and McDougall et al. (2012). *D*_*m*_ *>* 0 is the MMP-2 diffusion coefficient, and *Θ*_M_ *>* 0 and *Θ*_E/M_ *>* 0 are the rates of MMP-2 concentration provided by mesenchymal-like cancer cells and the partial-EMT cancer cells, respectively. Finally, *Λ >* 0 is the rate at which MMP-2 decays. Note that the mesenchymal-like and partial-EMT cancer cells also express MT1-MMP. However, MT1-MMP acts only locally where it is bound to the cancer cell membrane and its spatiotemporal evolution is hence congruent to that of the mesenchymal-like and of the partial-EMT cancer cells. Therefore, we do not include a separate equation.

Both the MT1-MMP expressed on the membranes of the mesenchymal-like and the partial-EMT cancer cells and the diffusive MMP-2 they secrete degrade the ECM. In the respective equation (3.2), this is expressed through the degradation rates *Γ*_M_ *>* 0 and *Γ*_E/M_ *>* 0 in the case of the MT1-MMP bound to the membranes of partial-EMT and of mesenchymal-like cancer cells, respectively, and for the diffusive MMP-2 by the degradation rate *Γ*_*m*_ *>* 0. Hence, given that we are disregarding ECM-remodelling for simplicity, the evolution of the ECM density 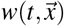 is governed by the following PDE:

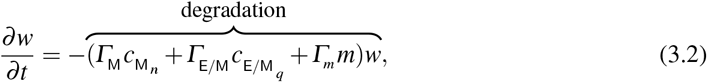

along with no-flux boundary conditions.

For the cancer cell migration on the grid we adopt a discrete approach where the movement probabilities of the cancer cells are given as follows:

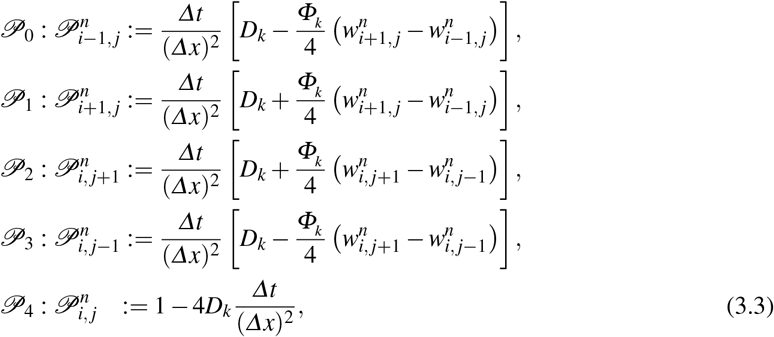

where *k* = E,E/M,M and, as throughout this paper, 0 *< D*_E_ *< D*_E/M_ *< D*_M_ and 0 = *Φ*_E_ *< Φ*_E/M_ *< Φ*_M_. 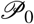, 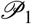, 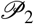, 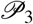 and 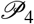 correspond to the probabilities that, during the next time step, a cancer cell at grid point (*x*_*i*_, *y*_*j*_) moves left, right, up, down, and not at all, respectively (Stéphanou et al., 2006; McDougall et al., 2012; Franssen et al., 2019a). Rules for proliferation and phenotypic transitions of the cancer cells (as well as—on the secondary grids—for cell death and dormancy) are then included as described below.

The more proliferative cancer cells of epithelial phenotype perform mitosis after time interval *T*_E_ and the less proliferative partial-EMT and mesenchymal-like cancer cells after time interval *T* and *T*_M_ (with *T*_E_ ≤ *T*_E/M_ ≤ *T*_M_), respectively. As previously in Franssen et al. (2019a), when proliferating, the cancer cells pass on their location so that a proliferating cancer cell is replaced by two daughter cells. Generally, during a proliferative step, cells are replaced by cells of their respective phenotype. However, in accordance with the biological findings presented in Section 2, the extended model allows for location-dependent full and partial EMT upon proliferation on the primary grid as follows:

- Cancer cells of epithelial phenotype may be replaced by a set of daughter cells consisting of one cell of epithelial and one of partial-EMT phenotype with probability 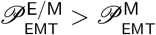 when proliferating;
- If at least one neighbouring grid point of a cancer cell of epithelial phenotype is unoccupied, it may be replaced by a set of daughter cells consisting of one of epithelial and one of partial-EMT phenotype with an additional probability 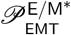;
- Cancer cells of epithelial and of partial-EMT phenotype may be replaced by a set of daughter cells consisting of one cell of epithelial and one of mesenchymal phenotype with probability 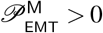 when proliferating.

As before, to account for competition for space and resources, the cancer cells on the respective grid point do not proliferate if there are already 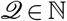 cancer cells on a grid point at the time of proliferation. Thus, 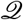 represents the preferred carrying capacity in our model.

### Intravasation

As in Franssen et al. (2019a), to represent the entry points into the blood vessels, a number of *U*_P_ ∈ ℕ_0_ normal blood vessels as well as *V*_P_ ∈ ℕ_0_ ruptured blood vessels are distributed throughout the primary grid. The normal blood vessels take the size of one grid point, while ruptured vessels consist of a group of *A*^*b*^ ∈ ℕ, where *b* = 1, 2,…, *V*_P_, adjacent grid points and can thus have different shapes. The entry rules for cancer cells of epithelial and mesenchymal phenotype remain as described in Franssen et al. (2019a). Moreover, in this extended framework, the cancer cells of partial-EMT phenotype are treated in the same way as those of mesenchymal phenotype in the sense that they may intravasate into both normal and ruptured vessels, unlike epithelial-like cancer cells.

### Travel through the vasculature

Cancer cells and cancer cell clusters remain in the vasculature for some time interval of length *T*_*V*_ ∈ ℕ, which biologically represents the average time the cancer cells spend in the blood system. Any cancer cells that enter a particular vessel at the same time are treated as one cluster and hence as a single entity once they are located in the vasculature. However, each cancer cell that is part of a cancer cell cluster disaggregates from its cluster with some probability 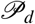 after spending a time interval of 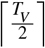 in the vasculature. After the time interval *T*_*V*_, the single cancer cells and the remaining cancer cell clusters are removed from the simulation unless they are randomly determined to survive. In accordance with the findings in Section 2, the survival probability is 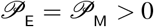 for single cancer cells of epithelial and mesenchymal phenotype, 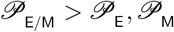 for single cancer cells of partial-EMT phenotype, and 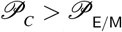 for cancer cell clusters.

### Metastatic growth

On the secondary grids 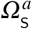, where *a* = 1, 2,…, *G*, the same phenotypes of cancer cells are accounted for as on the primary grid. Also, the same movement probabilities from equations (3.3) are used to describe their movement. However, we allow for organ-specific adjustment of the cell movement through differentiation of the respective diffusion and haptotactic coefficients, 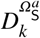 and 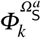, *k* = E,E/M,M, *a* = 1, 2,…, *G*.

Moreover, at the primary site, we modelled the assumption of well-adaptedness of the cancer cells to their tumour microenvironment of origin by considering proliferation every fixed time interval *T*_*k*_, *k* = E, E/M, M, if the carrying capacity *Q* permits. At the secondary sites, the cancer cells may not be as well-adapted to their new tumour microenvironment and may be exposed to the response of the immune system upon arrival. Furthermore, how well the cancer cells are adapted may vary between secondary organ tissues, *cf.* Section 2. To account for this, we make several adjustments on the secondary grids. Firstly, cancer cells may die with some grid-specific probability 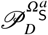 immediately prior to each potential proliferation. Secondly, a cell may not proliferate with some grid-specific probability 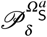 when proliferation is due to account for cancer cell dormancy. Besides this, if cancer cells do proliferate during a time step, we account for MET at the secondary sites in accordance with the biological findings presented in Section 2. Hence, cancer cells of partial-EMT phenotype on the secondary grids may be replaced by a set of daughter cells consisting of one cancer cell of epithelial and one of partial-EMT phenotype with probability 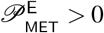 when proliferating. EMT does not occur on the secondary grids. However, as before, proliferation is capped as soon as a maximum of *Q* cancer cells per grid point is reached.

## 4. Setup of computational simulations and model calibration

To perform numerical simulations, we non-dimensionalised the system of equations (3.1)–(3.2) and the movement probabilities (3.3), with *k* = E, E/M, M, as described in Appendix A. As Anderson et al. (2000); Franssen et al. (2019a), we chose to rescale distance with an appropriate length scale *L* = 0.2 cm (since 0.1–1 cm is estimated to be the maximum invasion distance of cancer cells at an early stage of cancer invasion) and time with an appropriate scaling parameter 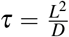. Here, *D* = 10^−6^ cm^2^s^−1^ is a reference chemical diffusion coefficient suggested by Bray (1992), such that *τ* = 4 × 10^4^ s, which corresponds to approximately 11 h.

We considered spatial domains of size [0, 1] × [0, 1]. This corresponds to physical domains of size [0, 0.2]cm × [0, 0.2]cm. In particular, we let the spatial domain *Ω*_P_ represent the primary site and the spatial domains 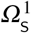, 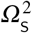 and 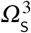 describe three potential metastatic sites. These spatial domains could represent *any* primary and secondary carcinoma sites. However, to give an example of a specific application, we chose *Ω*_P_ to represent the primary site of the breast, and 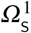, 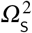 and 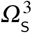 to correspond to the bones, lungs and liver, respectively, which are commonly observed metastatic sites in breast cancer.

The four spatial domains were discretised to contain 201 × 201 grid points each. This corresponds to a non-dimensionalised space step of *∆x* = *∆y* = 5 × 10^−3^, which results in a dimensional space step of 1 × 10^−3^ cm, and thus roughly corresponds to the diameter of a breast cancer cell (Vajtai, 2013). We then chose a time step of *∆t* = 1 × 10^−3^, corresponding to 40 s. This ensures that the scheme complies with the *Courant-Friedrichs-Lewy* (CFL) condition (Anderson et al., 2000), while still maintaining appropriate computational efficiency. We ran our simulation for a time period corresponding to ~ 24 days.

On each secondary grid, we chose 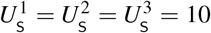 distinct grid points, on which blood vessels are located. For each grid, these blood vessels were placed randomly but at least two grid step widths away from the respective grid’s boundary. The same applies to the primary grid *Ω*_P_ but with the additional condition that the *U*_P_ = 8 single grid points, where normal blood vessels are located, and the *V*_P_ = 2 sets of five grid points, where ruptured blood vessels are placed, are located outside a quasi-circular region containing the 200 centre-most grid points. While these 10 randomly placed vessels are modelled to exist from the beginning, they represent those vessels that grow as a result of tumour-induced angiogenesis in the vascular tumour growth phase—hence they are placed away from the initial avascular epithelial tumour mass.

To represent a two-dimensional cross-section of a small avascular primary tumour, we placed a nodule that consisted of 288 randomly distributed epithelial-like cancer cells in the quasi-circular region of the 97 centre-most grid points of the primary grid. To account for competition for space, we allowed for no more than 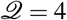 cancer cells on any grid point. This preferred carrying capacity of 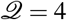 was applied throughout the simulation. The described initial condition ensured that the cancer cells were placed away from any pre-existing vessels to match the biology of an avascular tumour in epithelial tissue. Figure 7 gives an example of a typical initial cancer cell placement and vessel distribution on the primary grid.

**Figure 7.**
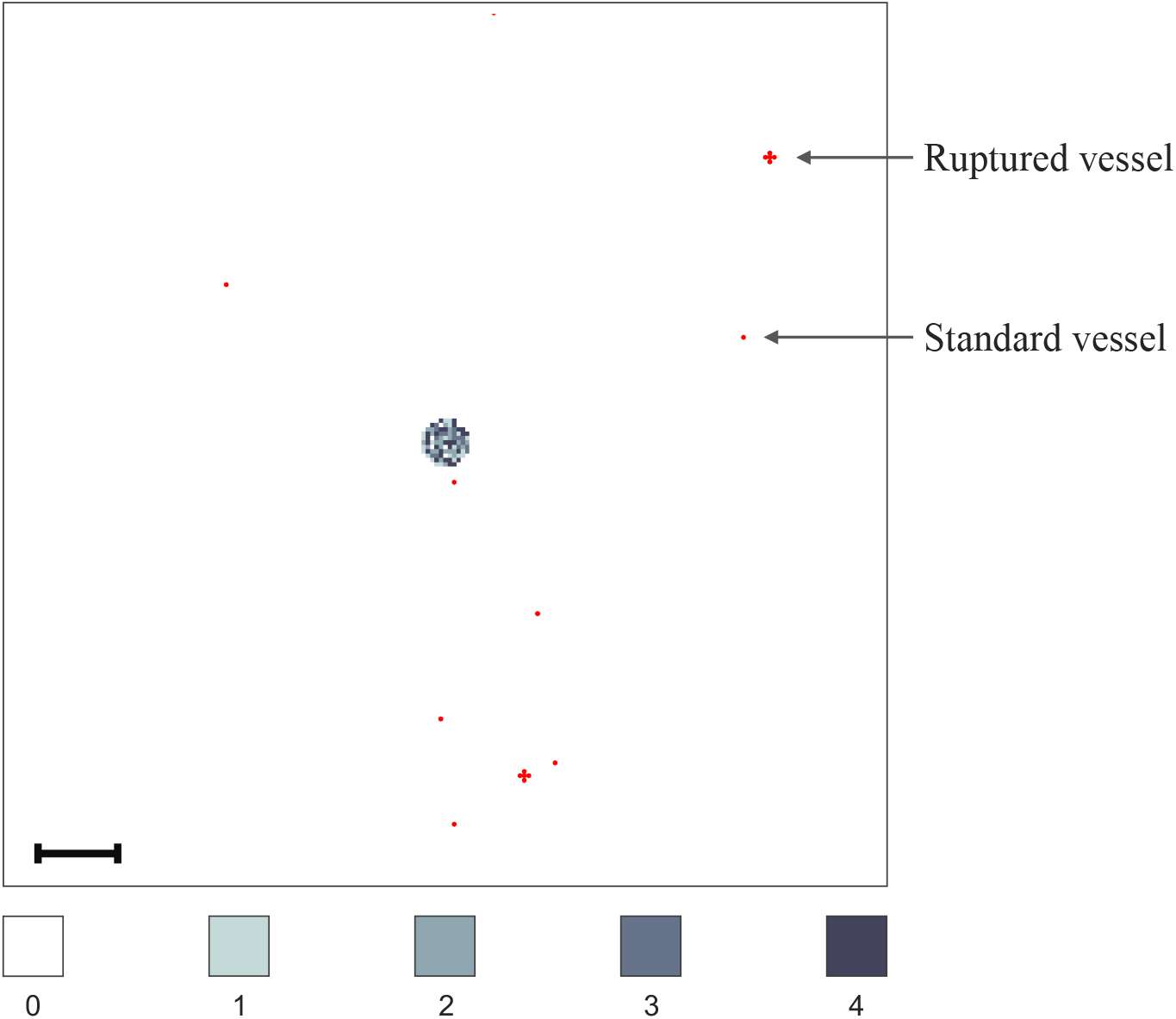
Vessel distribution and initial condition of cancer cells. The plot shows (in red) ten randomly distributed blood vessels on the primary grid, two of which are so-called ruptured vessels that consist of five rather than one grid point. In the centre of the grid, the initial distribution of epithelial-like cancer cells is shown. There are between 0 (white) and 4 (black) cancer cells on a grid point. As the initial distribution of cancer cells represents a 2D section through an avascular tumour, the blood vessels are placed at some distance away from the initial nodule of cancer cells. The scale bar denotes 0.02 cm.

In accordance with the ranges provided in Table A.1, we chose the epithelial-like cancer cell diffusion coefficient to be *D*_E_ = 1 × 10^−4^, the partial-EMT cancer cell diffusion coefficient to be *D*_E/M_ = 2.5 × 10^−4^ and set the mesenchymal-like cancer cell diffusion coefficient to *D*_M_ = 5 × 10^−4^. Furthermore, the epithelial, partial-EMT and mesenchymal haptotactic sensitivity coefficients were chosen to be *Φ*_E_ = 5 × 10^−5^, *Φ*_E/M_ = 1 × 10^−3^, and *Φ*_M_ = 2 × 10^−3^, respectively.

Taking into consideration the qualitative and quantitative biological findings in Section 2, we further assumed that, once in the vasculature, single CTCs of epithelial and mesenchymal phenotypes had a survival probability of 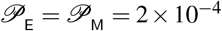, while those of partial-EMT phenotype survived the travel through the vasculature with probability 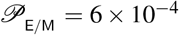. The survival probability of CTC clusters was set to 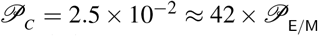, in accordance with the finding by Aceto et al. (2014) that the survival probability of CTC clusters is between 23 and 50 times higher than that of single CTCs. Surviving single CTCs and CTC clusters exited onto the secondary grids after spending a time period of *T*_*V*_ = 0.18 in the blood system, which corresponds to 2 h and hence to the breast cancer-specific clinical results in Meng et al. (2004).

Further, we assumed a uniform initial MMP-2 concentration of 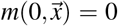 across all the spatial domains. We varied the initial ECM density according to the organ each grid represents using clinical measurements of ECM densities in organs from (ICRP, 2009). These are presented in Table A.1. For this, we took 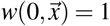, 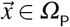, on the primary grid that represents the breast as our reference density. We then rescaled the initial ECM densities on the secondary grids relative to this initial density. For the bones, lungs and liver, respectively, this yielded 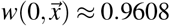, for 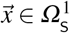, 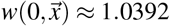, for 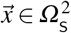, and 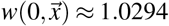, for 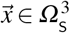. We assumed that epithelial-like cancer cells divide by mitosis every interval *T*_E_ = 2, the partial-EMT cancer cells every interval *T*_E/M_ = 3, and the mesenchymal-like cancer cells every *T*_E_ = 6. This corresponds to approximately 22 hours, 33 hours and 67 hours, respectively, which is consistent with the average doubling times found in breast cancer cell lines (Milo et al., 2009; NCI, 2015; Hughes et al., 2008). Moreover, we assumed that on the primary site, upon proliferation of a cancer cell of epithelial or partial-EMT phenotype, one of the daughter cells mutates into a mesenchymal-like cancer cell with probability 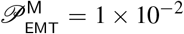. Similarly, one daughter cell of each epithelial-like cancer cell may mutate into a partial-EMT cancer cell with probability 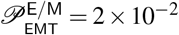 throughout the grid. Moreover, there is an additional probability from partial EMT of 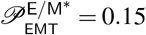 if the epithelial-like cancer cell is located at the edge of the tumour. Instead of representing the adaptation to each grid through these parameter settings, we determined the relative likelihood of metastasis-formation at the three secondary sites by consulting data on the transition probabilities of primary breast cancer to the metastatic sites of the bones, lungs and liver, respectively. As in Franssen et al. (2019a), we used data gathered in a study of 4181 breast cancer patients (Kuhn Laboratory, 2017). As shown in Figure 4 of Franssen et al. (2019a), the one-step transition probability from the breast to the bones was 23.1%, to the lungs was 15.3% and to the liver was 11.0%. Since we focus solely on the spread to these three metastatic sites and spread to other organs is included in the terms accounting for vascular death, we obtain the relative likelihoods of spread to the bones, lungs and liver, which are 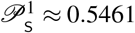, 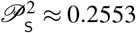, and 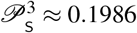, respectively.

At the secondary sites, cancer cells of partial-EMT phenotype were replaced by a set of daughter cells, consisting of one cell of epithelial and one of partial-EMT phenotype, with probability 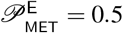 during proliferation. Due to a lack of organ-specific data on differences in the tumour microenvironments that could affect the diffusion and haptotactic coefficients of the cancer cells of various phenotypes—as well as their dormancy and death probabilities—at the time of writing, we restricted the differentiation between organs to their local initial ECM density at this stage. Accordingly, we took 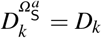 and 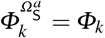, where *k* = E, E/M, M on all grids in our model. Similarly, the dormancy and death probabilities on all secondary sites were 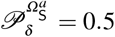 and 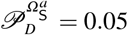, *a* = 1, 2, 3. An overview of the parameter values mentioned herein can be found in Table A.1.

## 5. Computational simulation results

To verify that our modelling framework is able to capture the key steps of the invasion-metastasis cascade, we ran simulations with the parameters shown in Table A.1. We provide sample results showing the primary and the three secondary grids at various times in the range of 0 to 24 days during one sample simulation in Figure 8 and Figures 9–11, respectively. We chose results on the primary grid to show the spatiotemporal dynamics on day 0, 11 and 22 so that they can be contrasted to those in Franssen et al. (2019a). For the secondary sites, we chose to present sample results for times that best give evidence of the various mechanisms related to metastatic spread, MET and the consequences of the immune response at secondary sites that are described through this modelling framework. However, this does not imply that these phenomena are limited to the times and locations depicted in Figures 8–11 in that particular or in other simulations.

**Figure 8.**
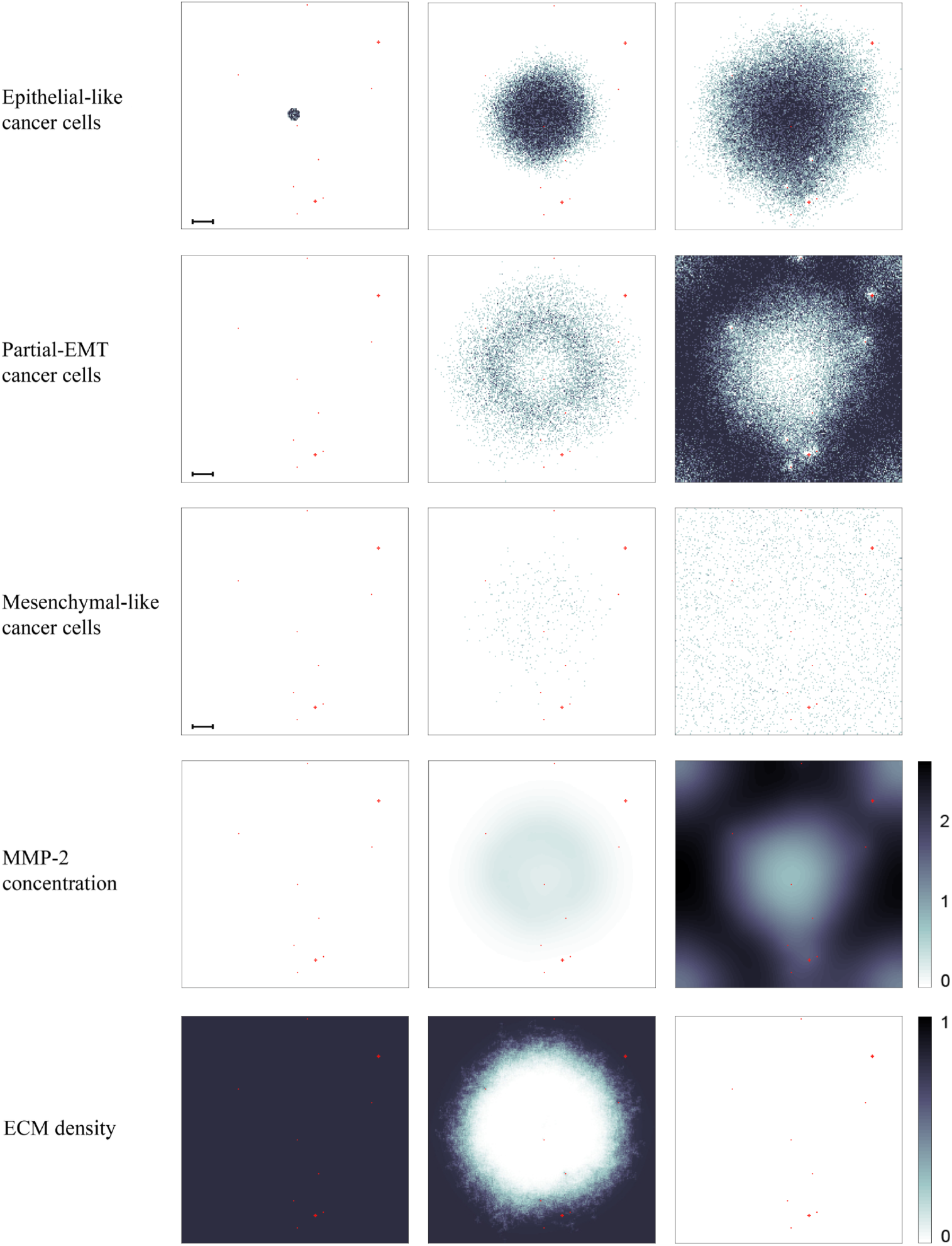
Simulation results on the primary grid. Primary tumour dynamics at 0 days, ~11 days and ~22 days. For each time step, the distribution of epithelial-like, partial-EMT and mesenchymal-like cancer cells (first to third row) is shown, with the discrete number of cancer cells per grid point ranging from 0 (white) to 4 (black) on each of the panels. The MMP-2 concentration (fourth row) continuously varies between 0 (white) and 2.6602 (black), and the ECM density (bottom row) between 0 and 1. Red dots represent blood vessels. There are 8 normal blood vessels of the size of one grid point as well as 2 ruptured blood vessels, which extend over 5 grid points each. If cancer cells are moved to these grid points, they may enter the vasculature and can potentially extravasate at secondary sites (*cf.* Figures 9–11). The scale bar denotes 0.02 cm and applies to all of the panels.

**Figure 9.**
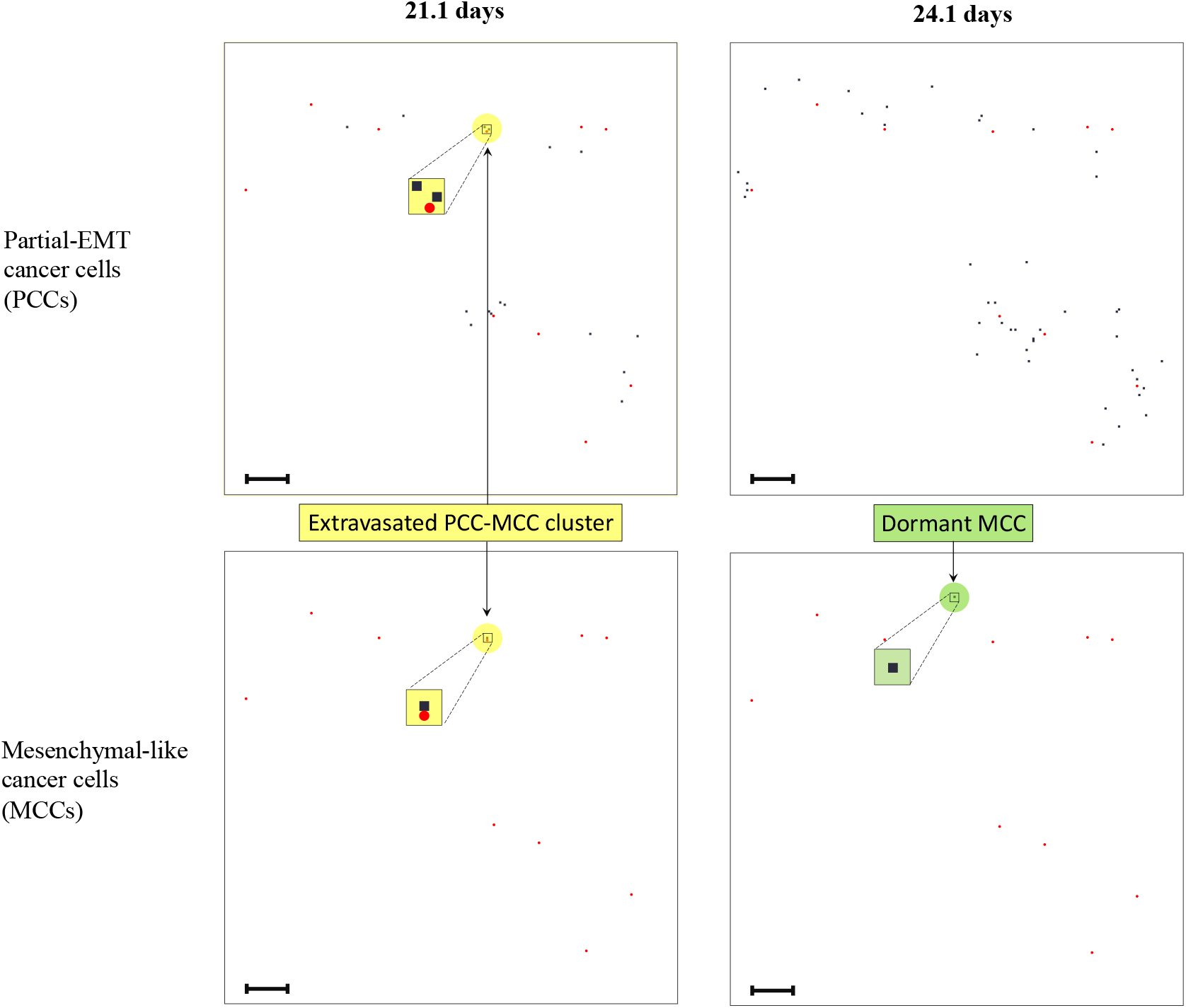
Cluster extravasation and dormancy on secondary grid representing the bones. Distribution of partial-EMT cancer cells (upper panels) and mesenchymal-like cancer cells (lower panels) at the secondary site representing the bones is shown after ~21 days (left) and ~23 days (right). The number of cancer cells per grid point varies between 0 (white) and 1 (black). Around day 21, a cluster consisting of two partial-EMT and one mesenchymal-like cancer cell extravasates onto the grid of the bones (yellow). Moreover, over the 3 day period between the panels on the left and on the right, the mesenchymal-like cancer cell, which normally has a doubling-time of ~2.78 days, remains dormant (green). The scale bar denotes 0.02 cm.

As described in Section 4, we started the simulations with a small nodule of epithelial-like cancer cells of diameter ~1.5 10^−2^ cm (*cf.* Figure 7). These were located on the primary grid representing the breast, which had an ECM of uniform density and contained no partial-EMT cancer cells, no mesenchymal-like cancer cells and no MMP-2, as shown in the left-most column of Figure 8. As the middle column of Figure 8 shows, after 11 days, the epithelial-like cancer cells had invaded the local tissue, covering a nearly circular area of approximately 0.1 cm diameter. Moreover, some partial-EMT and mesenchymal-like cancer cells could be observed on the primary grid. Their occurrence arose from cancer cells of previously epithelial-like phenotype via (partial) EMT. Both of these cell types occurred sparsely within a quasi-circular region with an approximate diameter of 0.18 cm. Additionally, the partial-EMT cancer cells populated a ring-shaped area at the edge of the tumour more densely. The MMP-2 concentration broadly followed the distribution of the partial-EMT cells and ranged from 0 to 0.38. Moreover, the ECM had been degraded in the centre of the grid and a density gradient could be observed at the edge of this near-circular region. After 22 days, the area occupied by the epithelial-like cancer cells in the centre of the tumour had expanded further. Also, the ring-like area populated with partial-EMT cancer cells at the tumour edge had grown and become more densely populated. The mesenchymal-like cancer cells were now sparsely spread throughout the whole grid. In general, we observed that areas on the grid near vessels were sparsely occupied, if at all. The distribution of the MMP-2 concentration still broadly followed the evolution of the partial-EMT cancer cells, now ranging from 0.76 to 2.66. The ECM on the domain that we considered had now been fully degraded.

In addition to the cancer cell invasion on the primary grid, we also observed metastatic spread to the grids representing the secondary sites. In Franssen et al. (2019a), we showed all of the secondary grids at the same time instances as the primary grid, so after approximately 11 and 22 days. Moreover, we included the spatiotemporal dynamics of the ECM density and of the MMP-2 concentration for each of the grids at these times. For these results, we hence refer the interested reader to this previous work. In this paper, we focus on the presentation of the additional phenomena captured through this extension of the model instead. As the newly introduced features are connected to the cells of various phenotypes that are included in the model, at the secondary sites we only show their evolution, while omitting the presentation of the MMP-2 and ECM dynamics. To give examples of how the various mechanisms that this model describes are instantiated in the simulations, we show the grids of the bones (Figure 9), the lungs (Figure 10) and the liver (Figure 11) at various times ranging from 16.3 to 24.1 days. The particular times differ between the secondary grids as they were chosen in order to best present how the phenomena occur in the simulations. Yet, within each grid, the time instances shown are such that the cell phenotypes depicted in the corresponding panels are at least the length of a cell doubling interval apart to allow all cells in the respective grid to have proliferated, if applicable, at least once.

**Figure 10.**
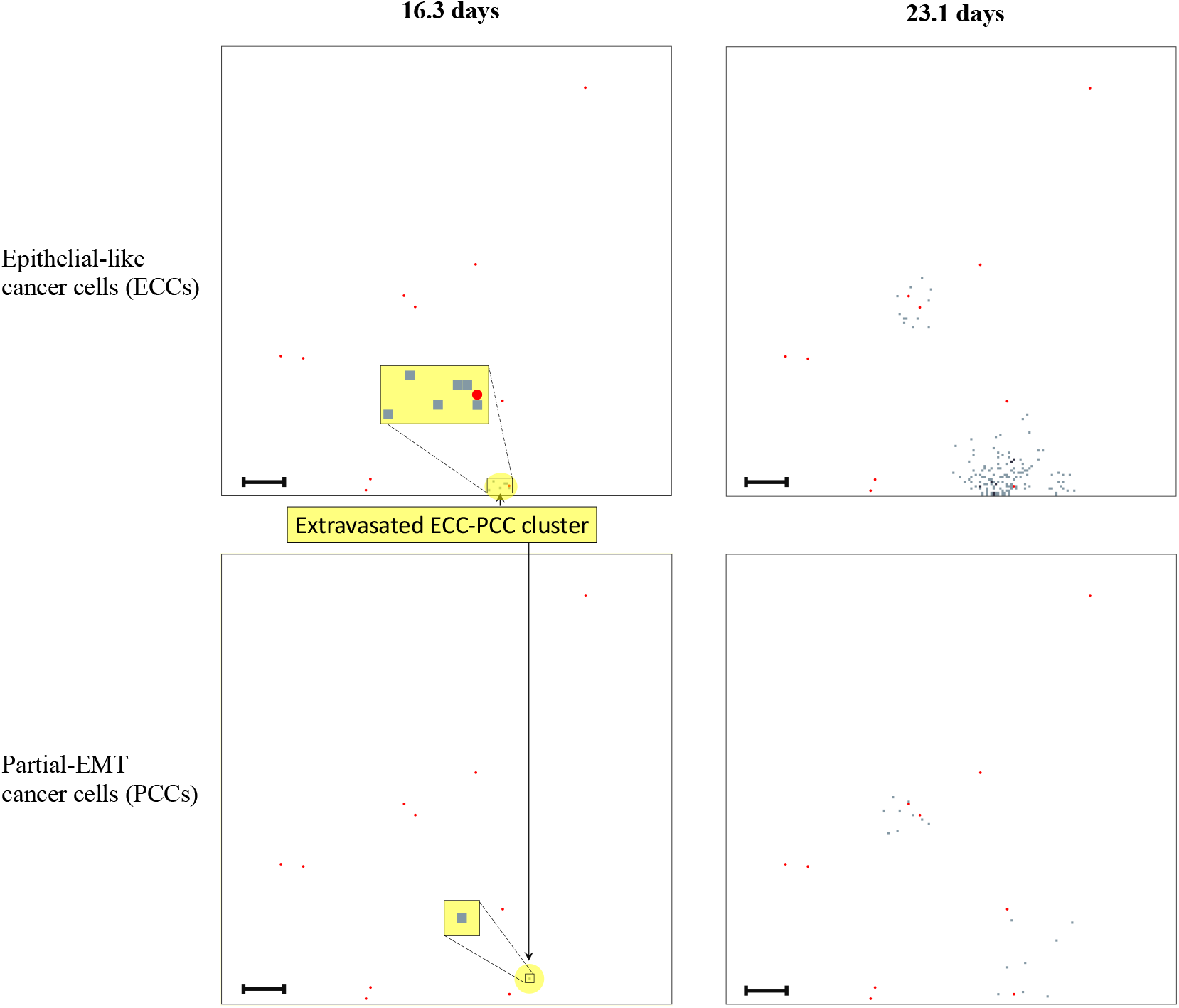
Cluster extravasation and largest metastatic lesion on secondary grid representing the lungs. Distribution of partial-EMT cancer cells (upper panels) and mesenchymal-like cancer cells (lower panels) at the secondary site representing the bones is shown after ~21 days (left) and ~23 days (right). The number of cancer cells per grid point varies between 0 (white) and 2 (black). Around day 21, a cluster consisting of six epithelial-like and one partial-EMT cancer cell extravasates onto the grid of the lungs (yellow). This early extravasation of a relatively large cluster of epithelial-like cancer cells results in the largest metastatic growth which can be observed in the right half of the panels on the right. The scale bar denotes 0.02 cm.

**Figure 11.**
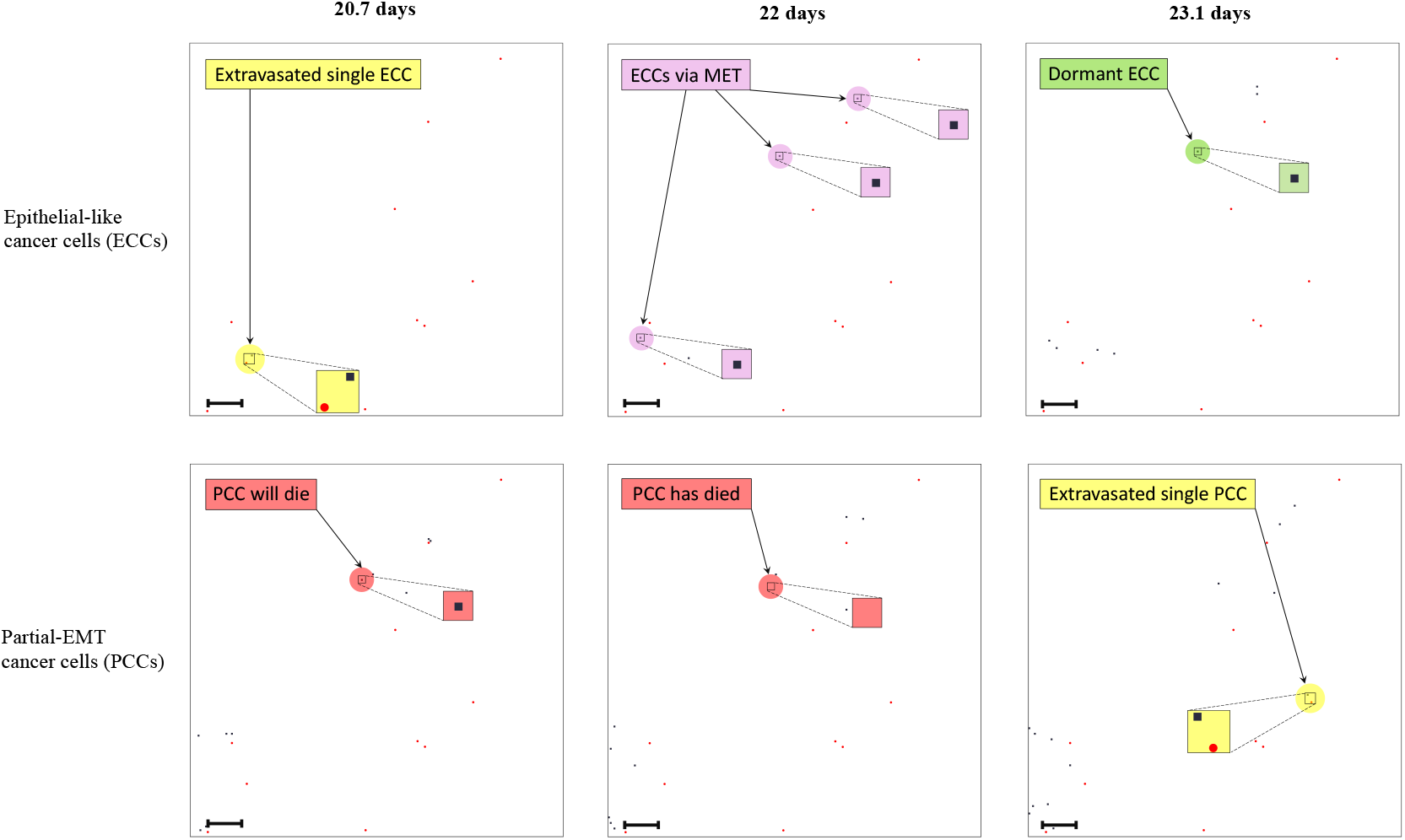
Single cell extravasations, MET, dormancy and cell death on secondary grid representing the liver. Distribution of partial-EMT cancer cells (upper panels) and mesenchymal-like cancer cells (lower panels) at the secondary site representing the bones is shown after ~21 days (left) and ~23 days (right). The number of cancer cells per grid point varies between 0 (white) and 1 (black). On day 20 and 23, a single epithelial-like and a single partial-EMT cancer cell extravasate onto the grid of the liver (yellow). No extravasations took place during the presented time period. Hence, the three epithelial-like cancer cells that occurred in the period between 20.7 days and 22 days in the upper middle panel (pink) are a result of MET of the partial-EMT cancer cells presented in the bottom row of panels. During the same time period, a partial-EMT cancer cell dies (red). Moreover, over the 1.1 day period between the panels in the middle and on the right, an epithelial-like cancer cell, which normally has a doubling-time of ~0.93 days, remains dormant (green). The scale bar denotes 0.02 cm.

We proceed by describing the results at the secondary sites grouped by the mechanisms that we aim to highlight (Figures 9–11) rather than grid-by-grid, as these mechanisms typically occur on all secondary grids. Furthermore, we present the dynamics of the cell-phenotype evolution of the population sizes on the secondary grids in Figures 12–13.

**Figure 12.**
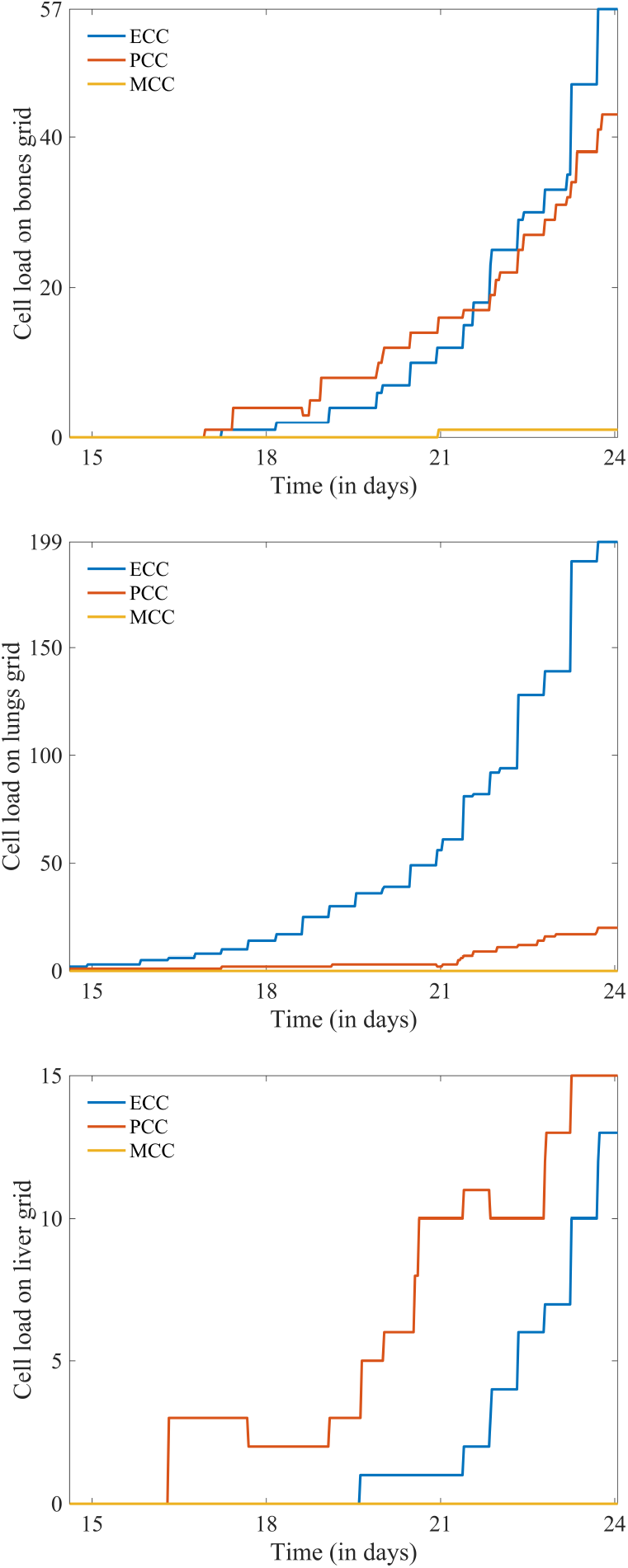
Phenotype-specific cell load over time on secondary grids. Plots of total number of epithelial-like (ECC; blue), partial-EMT (PCC; red) and mesenchymal-like (MCC; yellow) cancer cells on the grids of the bones, lungs and liver (top to bottom) in the period between 14.6 and 24.1 days. On each grid, the initial growth arises from an extravasation of cells. The stepwise, mostly non-negative growth pattern thereafter largely occurs from—often synchronous within the phenotypic group— proliferation of cells in combination with further extravasations. For ECCs, part of the growth also results from PCCs that undergo MET. As MET during PCC proliferation results in one PCC and one ECC, MET typically causes the PCC growth to slow down. Negative growth, as e.g. observed in the PCC population on the top ‘bones’ grid after day 18, on the middle ‘lungs’ grid at day 21, and on the bottom ‘liver’ grid after day 17 and 22, is always a result of cell death. Throughout, MCCs on the secondary grids remain rare.

### Extravasations

We observed extravasations of single cancer cells of various phenotypes, as well as of homogeneous and heterogeneous cancer cell clusters. Examples of a selection of these extravasations are highlighted in yellow on the grids representing the various secondary organs. Figure 11 shows samples of recently extravasated single cancer cells of epithelial and of partial-EMT phenotype on the grid representing the liver. Figures 9 and 10 show examples of extravasated cancer cell clusters consisting of mixed phenotypes. These consist of two partial-EMT and one mesenchymal-like cancer cell on the grid that represents the bones, and of six epithelial-like and one partial-EMT cancer cell in the case of the grid representing the lungs.

During the 22 day period over which the simulation on the primary grid was run, we observed 6 extravasations of single cells, as well as 11 of clusters consisting of two cells, 6 of clusters consisting of three cells and 1 extravasation each of clusters consisting of six and of seven cells. Another general observation was that during the simulated 24 day period, only one mesenchymal-like cancer cell successfully extravasated onto a secondary grid. All other extravasations were performed by single cancer cells as well as by homo- and heterogeneous cancer cell clusters, which were mainly of partial-EMT phenotype but also of epithelial-like phenotype. The highest number of extravasations of either a cancer cell or a cancer cell cluster was observed onto the grid of the bones.

### MET

On the grid representing the liver, no extravasations took place during the time period between 20.7 and 23.1 days, i.e. during the period shown in Figure 11. Hence, the three epithelial-like cancer cells that occurred during the period between 20.7 and 22 days, which are highlighted in pink in the upper row of panels, are a result of MET of the partial-EMT cancer cells presented in the bottom row of panels. If MET occurred during a proliferative step, the respective partial-EMT cancer cell was replaced by one cancer cell of its own phenotype as well as one of epithelial-like phenotype. Overall, the phenomenon of MET caused the growth rate of epithelial-like cancer cells to increase while slowing the growth rate of partial-EMT cancer cells at secondary sites. This trend is captured in the plots in Figure 13.

**Figure 13.**
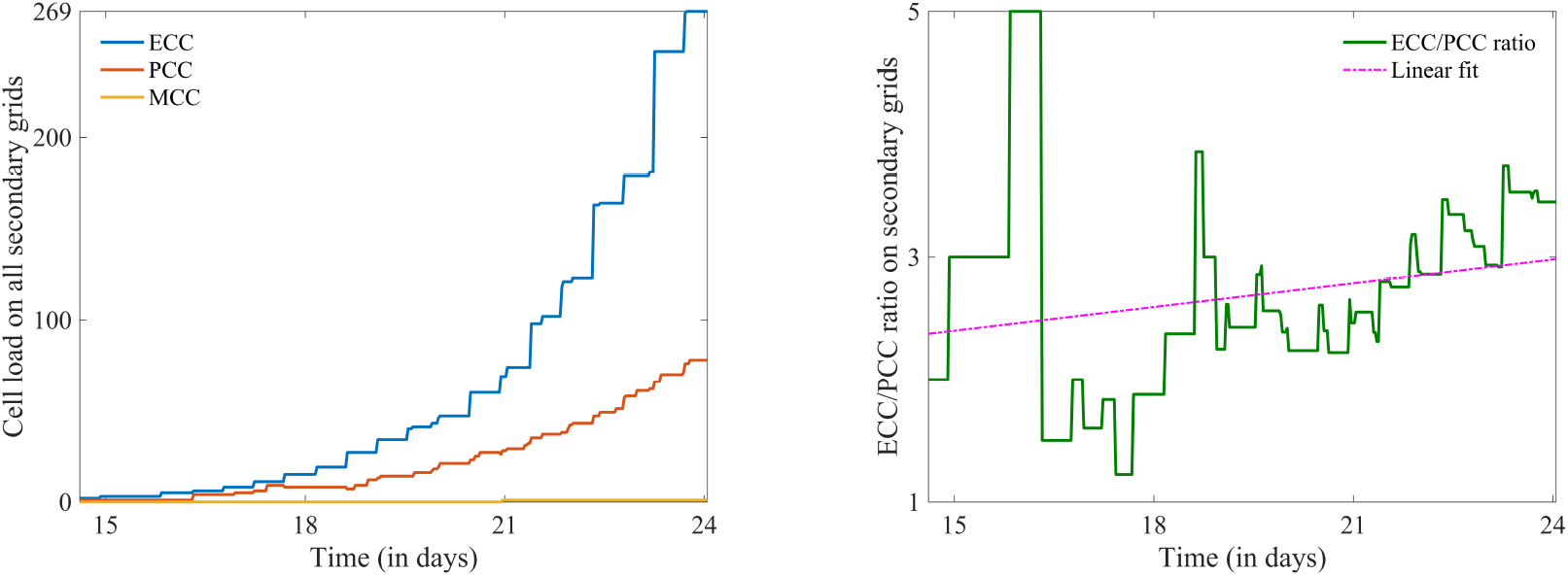
Trends in phenotype-specific cell load on secondary grids overall. Left: Combined total cell load on the secondary grids of cancer cells of epithelial-like (ECC; blue), partial-EMT (PCC; red) and mesenchymal-like (MCC; yellow) phenotype between the period of 14.6 and 24.1 days. The number of ECCs grows most rapidly over time—their growth is caused by extravasations, MET and proliferation. PCCs grow steadily but less rapidly. Their growth is slower due to their larger proliferation interval but also due to a subset of PCCs undergoing MET during proliferation. Only 1 MCC is observed over the time period. Right: Plot of the ratio of ECCs to PCCs over the same time frame (green). Throughout, there are more ECCs than PCCs and the ratio increases with time for the reasons explained above. The best linear fit line (pink) highlights this trend.

While any partial-EMT cancer cell can potentially undergo MET, the mesenchymal-like cancer cells cannot change phenotype. The sole mesenchymal-like cancer cell in Figure 9 is an example of such a phenotypically stable cell.

### Metastatic growth

The by far largest micrometastatic lesion during the simulation period presented itself on the grid of the lungs, shown in Figure 10. It resulted from a cluster consisting of six epithelial-like cancer cells and one partial-EMT cancer cell that extravasated relatively early—after approximately 16 days. All other lesions remained comparatively small during the same 23 day period, consisting of less than 20 cancer cells of almost exclusively epithelial-like and partial-EMT phenotypes. This is also reflected in the evolution of the total cell number on the three grids represented through the plots in Figure 12. As time progressed, a tendency towards a higher percentage of epithelial-like cancer cells at secondary sites was observed, as Figure 13 suggests.

### Dormancy

Given that we have chosen the periods between the time instances presented through the panels in Figures 9 to 11 to be such that there exists at least one opportunity for each cancer cell to reproduce, these figures show examples of dormant cancer cells at the secondary site of the bones and the liver, respectively. Due to their dormancy, the respective mesenchymal-like and epithelial-like cancer cells do not proliferate while other cells on the grids may. The two examples of dormant cells discussed in this section are highlighted in green in the respective figures.

### Death due to maladaptation & immune response

Figure 11 shows an example of a partial-EMT cancer cell that dies in the period between 20.7 and 22 days. Other examples of cell death on secondary sites become evident when examining the cell population growth plots for the partial-EMT cells on each of the secondary sites shown in Figure 12. Negative growth, as e.g. observed in the partial-EMT population on the top ‘bones’ grid after day 18, on the middle ‘lungs’ grid at day 21, and on the bottom ‘liver’ grid after day 17 and 22, is always a result of cell death. The fact that we only observe cell death in the partial-EMT population through these plots does not imply that cell death does cannot occur in the cell populations of other phenotypes. The epithelial-like cancer cells in the model tend to proliferate mostly synchronously. Hence, rare potential cell deaths are likely to be overshadowed in the plots in Figure 12 by an even larger positive cell growth at the same time instance. The same applies to potential other partial-EMT cell deaths.

## 6. Discussion and perspective

In this paper, we have extended the mathematical framework for the metastatic spread of cancer originally proposed in Franssen et al. (2019a) to include EMT and MET. As a result, the framework now additionally accounts for transitions of cancer cells between an epithelial, a newly introduced partial-EMT and a mesenchymal phenotypic state. This is achieved in a location-dependent fashion—both with respect to the steps of the invasion-metastasis cascade and with respect to the intra-tumoural location of the cancer cells. This way, the modelling framework captures the phenomena of EMT and MET in their physiological context. Furthermore, we include organ-specific differences in the local tissue of the secondary sites involved in our model by accounting for their ECM density in accordance with biological findings in (ICRP, 2009). Finally, the extended framework now also takes into account cancer cell dormancy and death as a result of maladaptation to the new tumour microenvironments at the secondary sites as well as due to the local immune response.

Through computational simulations, we found that the extended metastasis modelling framework provides biologically realistic outcomes and gives further insight into the above-described mechanisms that underpin the invasion-metastasis cascade at the cellular scale. Tumour shape and metastatic distribution at the primary site were predicted to appear as one would expect in a cancer patient who has not yet received treatment. In particular, we found that the partial-EMT cancer cells formed a ring-shaped leading front along the tumour edge, which was also seen in experiments (Nurmenniemi et al., 2009) as well as in human tissue, as shown in Figure 4 from Puram et al. (2017).

Nurmenniemi et al. (2009) further observed an average maximum invasion depth of 5.47 × 10^−2^ cm over 14 days when culturing HSC-3 cancer cells, a human oral squamous carcinoma cell line with high metastatic potential, on top of myoma tissue. This translates into an average maximum invasion speed of approximately 4.52 × 10^−8^ cm s^−1^. It suggests that our observed maximum invasion depth of ~9 × 10^−2^ cm in 11 days by partial-EMT and mesenchymal-like cancer cells and the resulting estimated average for the maximum invasion speed of approximately 9.38 × 10^−8^ cm s^−1^ are realistic results, given that migration speed varies between cancer cell lines and that the displacement of the cancer cells is likely a result of a combination of migration and proliferation.

The number of extravasations of single cancer cells or of a cancer cell cluster per secondary site further matched the clinical data of 4181 breast cancer patients, which underlie our simulations. The bones are the most frequently observed site of metastatic spread from primary breast cancer in the data processed by the Kuhn Laboratory (2017)—correspondingly, we observed the highest number of extravasations to the grid representing this site.

To our knowledge, there are currently no data available that claim to deliver an accurate estimation of the typical metastatic load from primary breast cancer to secondary sites over a specified time frame. However, we believe our results are biologically appropriate with regards to their timings. They are in correspondence with the conclusion reached by Obenauf & Massagué (2015) in their review of the metastatic traits that allow cancer cells to colonise various secondary sites, suggesting that CTCs and metastatic spread can be detected soon after vascularisation of the primary tumour, as in our simulations.

The types of extravasations that we observed through the simulations in our model coincide with the biological evidence presented in Section 2 that CTCs of all phenotypes appear to be able to extravasate (Banyard & Bielenberg, 2015). Furthermore, only a low proportion of extravasations included mesenchymal-like cancer cells—the bulk of extravasating cells were of partial-EMT phenotype and others of epithelial phenotype.

As discussed above, the highest number of extravasations was observed onto the grid of the bones. Yet, as Figure 10 indicates, the largest micrometastasis, which resulted from the early metastatic spread of a large cluster consisting predominantly of epithelial-like cancer cells, occurred at the site of the lungs, where only two extravasations were observed over the total time period that we considered. This emphasises that cancerous spread is highly complex and difficult to predict, a feature represented through the stochasticity involved in multiple processes of our model. Examples of such processes are (partial) EMT at the primary site, the survival of CTCs and the potential partial or full dissemination of CTC clusters in the vasculature, the determination of the secondary site of extravasation, as well as MET, dormancy and cell death at secondary sites. Furthermore, the fact that the largest growth stemmed from a cluster consisting of predominantly epithelial-like cancer cells highlights that this cell type with its distinguishing feature of rapid proliferation is generally the one best adapted to growth in the tumour microenvironment at secondary sites. This observation and our observation that—as time progresses— increasing numbers of partial-EMT cancer cells transit to an epithelial-like phenotype coincide with two of the biological findings discussed in Section 2. The first such finding is that the bulk of cancer cells at secondary sites are of epithelial-like phenotype (Pastushenko & Blanpain, 2018) as well as some of partial-EMT phenotype (Dongre & Weinberg, 2019). The second finding in agreement with our results is the observation by Ruscetti et al. (2015) that macrometastases at the secondary site of the lungs consisted mainly of epithelial-like cancer cells while smaller lesions presented few epithelial-like cancer cells and thus mainly cells with some degree of mesenchymal-traits. Finally, our model accounts for the biological evidence presented in Ocaña et al. (2012); Kröger et al. (2019) that cancer cells of a stable mesenchymal-like phenotype are unable to transform via MET and hence fail to give rise to metastatic growth at secondary sites.

In the current modelling approach, we account for the fact that EMT and MET have been observed to occur in specific steps of the invasion-metastasis cascade as well as in specific locations within the primary tumour. For instance, partial EMT appears to be triggered predominantly at the primary site and towards the tumour boundary, as observed *in situ*—see Figure 4 and Puram et al. (2017). Also, in the early stages of colonisation at a secondary site, MET is the predominant mutation. It would be desirable to additionally include a *physiological* motivation for the mutations that we model, like e.g. developed in Sfakianakis et al. (2017). In particular, we aim to incorporate the physiological motivation by accounting for the role of hypoxia as a trigger for EMT and MET in the following sense. While the full spectrum of mechanisms underlying the induction of EMT remains elusive to date (Wang et al., 2016), it is assumed that tumour-induced hypoxia plays an important role in the process (Imai et al., 2003; Yang et al., 2008; Wang et al., 2016; Petrova et al., 2018). The hypoxic environment in the tumour activates its main effector *hypoxia-inducible factor-1* (HIF-1) (Petrova et al., 2018), which in turn activates EMT-TFs like Snail and Twist (Imai et al., 2003; Yang et al., 2008), thus promoting EMT and metastatic phenotypes. A biological model that connects the occurrence of tumour-induced hypoxia with EMT and angiogenesis via CAFs has recently been proposed in Petrova et al. (2018). The hypothesis is made that rapid tumour growth, which reduces the oxygen concentration in tumour and stroma regions far away from vessels since the diffusion of oxygen is limited to 100–200 µm, creates hypoxic regions. Epithelial-like cancer cells in these hypoxic regions produce signalling molecules that transform normal fibroblasts as well as other healthy cells in the stroma to CAFs (Zeisberg et al., 2007; Petrova et al., 2018). These CAFs have been shown to produce stiff aligned ECM. This differently organised ECM is, in turn, hypothesised to induce EMT in premalignant epithelial cells and to support cell migration in breast cancer (Dumont et al., 2013). CAFs have further been shown to promote angiogenesis via the production of vascular endothelial growth factor-C (VEGF), C-X-C motif chemokine 12 (CXCL12) and basic fibroblast growth factor (FGF-2) (Pietras & Östman, 2010), making hypoxia an angiogenic stimulus (Carmeliet & Jain, 2000). Our individual-based spatial modelling framework meets the prerequisites for an extension that includes the above-described biological phenomena. Therefore, we will connect the EMT features currently included in the metastasis framework with the prevalence of tumour-induced acutely and chronically hypoxic regions as well as with angiogenesis in future work.

To create an organ-specific model, we have taken into account differences in the local tumour microenvironment of primary and secondary organs in the body in two ways. Firstly, we aligned the relative likelihood of successful secondary spread to the organs in our model to the metastatic transition probabilities of breast cancer from large patient studies (Kuhn Laboratory, 2017). Secondly, we distinguished between the relative differences in ECM density between the organs according to biological measurements in (ICRP, 2009). We are aware that the variations in ECM density have a minimal influence on the cell movement at the moment, given that their movement is dependent on the ECM gradient rather than the absolute ECM density. However, we are looking to include biomechanical properties such as the ECM stiffness in the model, which has been shown to influence cell behaviour through the activation of intracellular signalling pathways (Kalli & Stylianopoulos, 2018). At this point, organ-specific and intra-organic differences in ECM will have a larger impact. Also, accounting for organ-specific differences in these two ways is, of course, a simplification of the actual physiology in many ways. For instance, in reality, differences between organs are not limited to the relative densities of their ECM. As explained in detail in Barney et al. (2016), the tissue-specific differences in the tumour microenvironment found in the organs are manifold and only marginally established. They include, for instance, the genetic markers associated with tissue-specific metastasis, the healthy cells typically found in these tissues, the ECM stiffness and its protein composition, and the tissue dimensionality. Also, the tumour microenvironments have been shown to evolve with time, resulting in e.g. pre-metastatic niche formation that has been observed both in mouse models and clinical studies (McAllister & Weinberg, 2014). Further, these and other features will not only differ between organs but also when considering the same organ in any number of patients. For this reason, it is our goal to include the metastatic programmes of the various organs in our model, once more is known about them. Until then, we will continue using the transition probabilities from large studies as Disibio & French (2008); Kuhn Laboratory (2017) to differentiate between the relative success of metastatic spread to the various organs.

Additional ideas on how to further develop this modelling framework are elaborated in Franssen et al. (2019a). These include extending the model to a third spatial dimension as well as accounting for biomechanical properties, re-seeding, pre-metastatic niche formation, and immune system activation. Moreover, treatment regimes could be modelled. One example is the inclusion of chemotherapy in the framework, especially once we account for angiogenesis, in a similar way to Powathil et al. (2012, 2013). Other approaches to modelling chemo-, radio-, nano- and immunotherapy, as well as targeted, hormone and combination therapy, some of which could function as a basis to modelling treatment approaches using this framework, have recently been reviewed by Chamseddine & Rejniak (2019).

## Appendices

## A. Previously established metastasis modelling framework

The general spatial modelling framework of the metastatic spread of cancer developed in Franssen et al. (2019a) is appended here for ease of accessibility. This previous modelling framework underlies the changes to the model introduced in the main body of this paper in Section 3. These changes have the purpose of including EMT, MET as well as the third partial-EMT cancer cell phenotype into the existing framework. Further changes are the differentiation of organ tissue via an organ-specific initial ECM density as well as the inclusion of cell death and dormancy at secondary sites in the body.

To account for cancer cell metastasis in a spatially explicit manner, we consider *G* + 1 spatial domains. These consist of the spatial domain representing the primary tumour site, *Ω*_P_ ⊂ ℝ^2^, as well as the *G* ∈ ℕ spatial domains representing the sites of potential secondary metastatic spread, 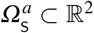, where *a* = 1, 2,…, *G*. In these spatial domains, we represent the MMP-2 concentration and the ECM density at positions 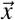 at time *t* by the continuous functions 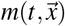 and 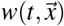, respectively, while capturing the spatiotemporal evolution of epithelial-like and mesenchymal-like cancer cells *individually*. We model the local cancer cell invasion by expanding the modelling approach used in Anderson & Chaplain (1998); Anderson et al. (2000) to our specific biological problem. However, we include a second cancer cell phenotype and also additionally consider MT1-MMP, which is taken to be bound to the membranes of the mesenchymal-like cancer cells and thus follows their discrete spatiotemporal dynamics. We designate locations in the primary spatial domain to function as entry points into the vasculature and, similarly, impose a spatial map of exit locations from the vasculature onto the secondary metastatic domains. This allows cancer cells to travel from the primary tumour site to secondary sites via blood vessels.

We next consider one key step of the invasion-metastasis cascade after the other. To make the key steps more recognisable, we begin each paragraph by printing the description of the corresponding step in the invasion-metastasis cascade (*cf.* Section 2 in Franssen et al. (2019a)) in bold. Further, the same step descriptions can be found on the left of the flowchart in Figure 5 in Franssen et al. (2019a). This highlights which parts of our model correspond to which sections in the text.

### Local cancer cell invasion

We adopt a discrete cell approach where the movement of cancer cells from grid point to grid point is accounted for by movement probabilities consisting of an unbiased component (*cf.* random motion) and a biased component proportional to gradients in the ECM (*cf.* haptotaxis). This way, we obtain the movement probabilities of the *individual* epithelial-like and mesenchymal-like cancer cells in equation (3.3) with *k* = E, M. Modelling the cancer cells individually allows us to track the evolution of single epithelial-like and mesenchymal-like cancer cells with different phenotypes, as well as their evolution.

The model we have described so far accounts for the movement of the cancer cells only. We thus need to additionally account for the *proliferation* of cancer cells in our model. The two cancer cell types included in our model proliferate at different frequencies. The more proliferative epithelial-like cancer cells perform mitosis after time interval *T*_E_, the less proliferative mesenchymal-like cell types after *T*_M_ (with *T*_M_ *> T*_E_). When proliferating, the cancer cells pass on their respective phenotype as well as their location so that a proliferating cancer cell is replaced by two daughter cells after the proliferative step has been performed. However, to account for competition for space and resources, the cancer cells on the respective grid point do not proliferate if there are 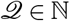 cancer cells on a grid point at the time of proliferation.

With reference to the flowchart shown in Figure 5 in Franssen et al. (2019a), the part of our approach described so far corresponds to *Movement, EMT & cell proliferation*, depicted in the upper region of the flowchart.

The mesenchymal-like cancer cells in our model have the ability to express diffusible MMP-2. The MMP-2 concentration 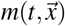 hence develops according to the equation:

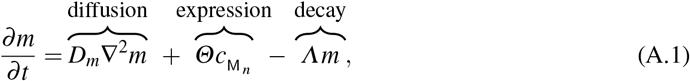

along with zero-flux boundary conditions. Here, 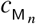, where *n* = 0, 1, 2, 3, 4, denotes the respective integer numbers of cells between 0 and 4 to indicate the presence of up to four mesenchymal-like cancer cells at a given position (*cf.* Stéphanou et al. (2006); McDougall et al. (2012)), *D*_*m*_ *>* 0 is the constant MMP-2 diffusion coefficient, *Θ >* 0 is the constant rate of MMP-2 concentration provided by mesenchymal-like cancer cells, and *Λ >* 0 is the constant rate at which MMP-2 decays. Note that the mesenchymal-like cancer cells also express MT1-MMP. However, MT1-MMP acts locally only where it is bound to the cancer cell membrane and its spatiotemporal evolution is hence congruent to that of the mesenchymal-like cancer cells. Therefore, we do not include a separate equation.

The diffusible MMP-2 degrades the ECM with a degradation rate of *Γ*_2_ *>* 0. The MT1-MMP expressed on the membrane of the mesenchymal-like cancer cells also degrades the ECM, which is expressed through the degradation rate *Γ*_1_ *>* 0. Hence, given that we are disregarding ECM-remodelling for simplicity, the evolution of the ECM density 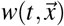 is governed by the following PDE:

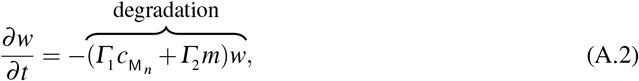

along with zero-flux boundary conditions. As before, 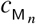, where *n* = 0, 1, 2, 3, 4, denotes the respective integer numbers of cells between 0 and 4 to indicate the presence of up to four mesenchymal-like cancer cells at a given position.

Since the continuous evolution of the MMP-2 concentration and of the ECM density is governed by equations (A.1) and (A.2), while the spatiotemporal evolution of the cancer cells (and, intrinsically, of the membrane-bound MT1-MMP) is captured by an individual-based model (*cf.* equation (3.3)), we model cancer cell invasion in a hybrid-discrete continuum approach of the kind pioneered by Anderson & Chaplain (1998) in their tumour-angiogenesis model. This approach was subsequently also used to model tissue invasion by cancer cells (Anderson et al., 2000; Anderson, 2005) and spatial evolutionary games (Burgess et al., 2016, 2017).

### Intravasation

With the model setup we have described so far, the cancer cells can invade the tissue locally in the primary spatial domain but cannot reach the spatially separated secondary domains. To allow for metastatic spread, we account for the connection of the primary spatial domain to the secondary spatial domains by incorporating blood vessels in our modelling framework. Examples of primary and secondary domains are presented in Figure 6 of Franssen et al. (2019a). To represent the entry points into the blood vessels, a number of *U*_P_ ∈ ℕ_0_ normal blood vessels as well as *V*_P_ ∈ ℕ_0_ ruptured blood vessels are distributed on the primary grid. The normal blood vessels take the size of one grid point, while ruptured vessels consist of a group of *A*^*b*^ ∈ ℕ, where *b* = 1, 2,…, *V*_P_, adjacent grid points and can thus have different shapes. Each secondary grid 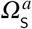 also has, respectively, 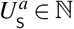 normal blood vessels, where *a* = 1, 2,…, *G* as before, that take the form of a single grid point each. On the primary grid, the grid points where the vessels are located allow the cancer cells to intravasate, while the respective grid points on the secondary grid allow for extravasation.

If, by the movement submodel described in Appendix B of Franssen et al. (2019a), a cancer cell on the primary grid is placed on a grid point that represents a blood vessel, it *may* leave the grid and enter the vasculature. Whether or not a cancer cell can successfully intravasate depends both on its own phenotype and on the type of vessel it is placed on.

Whenever a mesenchymal-like cancer cell is moved to a grid point (*x*_*i*_, *y*_*j*_) ∈ *Ω*_P_^1^, on which a *normal* single blood vessel is located, it will successfully enter the vasculature. Further, to represent collective invasion in the form of co-presence of mesenchymal-like and epithelial-like cancer cells, cancer cells of any type on the four neighbouring primary grid points (*x*_*i*+1_, *y*_*j*_), (*x*_*i*−1_, *y*_*j*_), (*x*_*i*_, *y*_*j*+1_) and (*x*_*i*_, *y*_*j*−1_) are forced into the vasculature together with the mesenchymal-like cancer cell on (*x*_*i*_, *y*_*j*_). Hence, a mesenchymal-like cancer cell moving to a grid point on which a normal blood vessel is located results in either a single mesenchymal-like cancer cell or a cluster consisting of up to 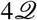 cancer cells of any phenotype intravasating. However, if an epithelial-like cancer cell is moved to a grid point (*x*_*i*_, *y*_*j*_) ∈ *Ω*_P_ where a *normal* single vessel is located without a mesenchymal-like cell being present there, the epithelial-like cancer cell will not intravasate and the grid point (*x*_*i*_, *y*_*j*_) will be treated like any other grid point. This is to model the fact that epithelial-like cancer cells have been shown to be unable to actively intravasate on their own.

Further, a cancer cell on the primary grid can move to one of the grid points where a *ruptured* vessel is located. Contrary to the above-described scenario of entering a normal vessel, a cancer cell of any type, which is placed on a grid point representing part of a ruptured vessel, can enter the circulation. The respective cancer cell takes with it any other cancer cells residing both on the grid points representing the ruptured blood vessel and on the regular grid points bordering the ruptured vessel. Biologically, the fact that cancer cells of any phenotype can intravasate mirrors that these blood vessels are already ruptured due to trauma or pressure applied by the expanding tumour, making the requirement of MDE-mediated degradation of the vessel wall redundant. The fact that other cancer cells on bordering grid points will enter the circulation together with cancer cells placed on grid points representing blood vessels captures some degree of the cell-cell adhesion found in collectively invading cancer cell clusters.

### Travel through the vasculature

If a cancer cell of either phenotype or a cluster of cancer cells successfully enters the vasculature either through a ruptured or a normal vessel, it will be removed from the primary grid and moved to the vasculature. Cancer cells and cancer cell clusters remain in the vasculature for some time *T*_*V*_ ∈ ℕ, which biologically represents the average time the cancer cells spend in the blood system. Any cancer cells that enter a particular vessel at the same time are treated as one cluster and hence as a single entity once they are located in the vasculature. However, each cancer cell that is part of a cancer cell cluster disaggregates from its cluster with some probability 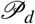 after 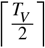 time steps. At the end of the time period *T*_*V*_, the single cancer cells and the remaining cancer cell clusters are removed from the simulation unless they are randomly determined to survive. The survival probability is 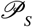 for single cancer cells and 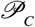 for cancer cell clusters.

### Extravasation

Any surviving cancer cells and cancer cell clusters are placed on one of the *G* secondary grids 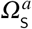 with probability ℰ_1_, ℰ_2_,…, ℰ_*G*_, where 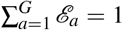. Also, on each specific secondary grid, the cancer cells extravasate through one of the randomly chosen 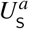 grid points that represent a blood vessel with equal probability. If the respective grid point cannot accommodate all of the entering cancer cells without violating the carrying capacity 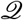, the remaining cancer cells are randomly distributed onto the four non-diagonally neighbouring grid points until these are filled to carrying capacity 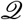. If there are further cancer cells to be placed onto the respective grid point at this instance, such cancer cells are killed to capture the influence of competition for space in combination with vascular flow dynamics.

### Metastatic growth

If and when cancer cells reach a secondary grid, they behave (i.e. replicate, move, produce MDEs etc.) there according to the same rules as on the primary grid, as indicated on the bottom of the flowchart in Figure 5 of Franssen et al. (2019a).

## B. Parameter settings used in the simulations

**Table A.1:** Parameter settings used in the simulations. In the first column, non-dimensional parameters are indicated by upper-case notation. Corresponding dimensional parameters are stated in brackets using lower-case notation. In the fourth column, we reference other mathematical modelling papers in brackets and biological papers without brackets. Epithelial-like, partial-EMT and mesenchymal-like cancer cells are represented by the acronyms ECC, PCC and MCC, respectively.

|  | Description | Non-dimensional value | Biological reference (Modelling reference) | Original value from cited source |
| --- | --- | --- | --- | --- |
| $\Delta t$ | Time step | $1 \times 10^{-3}$ | | 40 s |
| $\Delta x,$<br>$\Delta y$ | Space step | $5 \times 10^{-3}$ | Breast cell diameter in Vajtai (2013) | $1 \times 10^{-3}$ cm |
| $D_E$<br>($d_E$) | ECC diffusion coefficient | $1 \times 10^{-4}$ | Bray (1992)<br>(Anderson & Chaplain (1998))<br>(Deakin & Chaplain (2013)) | $1 \times 10^{-10}$ cm<sup>2</sup>s<sup>-1</sup> |
| $D_{E/M}$<br>($d_{E/M}$) | PCC diffusion coefficient | $2.5 \times 10^{-4}$ | Bray (1992)<br>(Anderson & Chaplain (1998))<br>(Deakin & Chaplain (2013)) | $1 \times 10^{-10}$ cm<sup>2</sup>s<sup>-1</sup> |
| $D_M$<br>($d_M$) | MCC diffusion coefficient | $5 \times 10^{-4}$ | Bray (1992)<br>(Anderson & Chaplain (1998))<br>(Deakin & Chaplain (2013)) | $1 \times 10^{-10}$ cm<sup>2</sup>s<sup>-1</sup> |
| $\Phi_E$<br>($\phi_E$) | ECC haptotactic sensitivity coefficient | $5 \times 10^{-5}$ | Stokes et al. (1990)<br>(Anderson & Chaplain (1998)) | $2.6 \times 10^3$ cm<sup>2</sup>M<sup>-1</sup>s<sup>-1</sup> |
| $\Phi_{E/M}$<br>($\phi_{E/M}$) | PCC haptotactic sensitivity coefficient | $1 \times 10^{-3}$ | Stokes et al. (1990)<br>(Anderson & Chaplain (1998)) | $2.6 \times 10^3$ cm<sup>2</sup>M<sup>-1</sup>s<sup>-1</sup> |
| $\Phi_M$<br>($\phi_M$) | MCC haptotactic sensitivity coefficient | $2 \times 10^{-3}$ | Stokes et al. (1990)<br>(Anderson & Chaplain (1998)) | $2.6 \times 10^3$ cm<sup>2</sup>M<sup>-1</sup>s<sup>-1</sup> |
| $D_m$<br>($d_m$) | MMP-2 diffusion coefficient | $1 \times 10^{-3}$ | Collier et al. (2011) | $1 \times 10^{-9}$ cm<sup>2</sup>s<sup>-1</sup> |
| $\Theta_E$<br>($\theta_E$) | MMP-2 production rate by ECCs | 0 | Estimated | |
| $\Theta_{E/M}$<br>($\theta_{E/M}$) | MMP-2 production rate by PCCs | 0.1 | Estimated | |
| $\Theta_M$<br>($\theta_M$) | MMP-2 production rate by MCCs | 0.195 | Estimated | |
| $\Lambda$<br>($\lambda$) | MMP-2 decay rate | 0.1 | Estimated in<br>(Deakin & Chaplain, 2013) | $2.5 \times 10^{-6}$ s<sup>-1</sup> |
| $\Gamma_1$<br>($\gamma_1$) | ECM degradation rate by MT1-MMP | 1 | Based on<br>(Deakin & Chaplain, 2013) | $1 \times 10^{-4}$ s<sup>-1</sup> |
| $\Gamma_2$<br>($\gamma_2$) | ECM degradation rate by MMP-2 | 1 | Based on<br>(Anderson et al., 2000) | $1 \times 10^{-4}$ M<sup>-1</sup>s<sup>-1</sup> |
| $T_V$ | Time CTCs spend in the vasculature | 0.18 | Meng et al. (2004) | $7.2 \times 10^3 \text{ s}$ |
| $T_E$ | ECC doubling time | 2 | Milo et al. (2009); NCI (2015) | $8 \times 10^4 \text{ s}$ |
| $T_{E/M}$ | PCC doubling time | 3 | Milo et al. (2009); NCI (2015) | $1.2 \times 10^5 \text{ s}$ |
| $T_M$ | MCC doubling time | 6 | Milo et al. (2009); NCI (2015)<br>Hughes et al. (2008) | $2.4 \times 10^5 \text{ s}$ |
| $\mathcal{P}_\delta^{\Omega_S^{1,2,3}}$ | Probability no cell proliferation when due on $\Omega_S^{1,2,3}$ | 0.5 | Estimated | |
| $\mathcal{P}_D^{\Omega_S^{1,2,3}}$ | Probability cell death before proliferation on $\Omega_S^{1,2,3}$ | 0.05 | Estimated | |
| $\mathcal{P}_E = \mathcal{P}_M$ | Epithelial/mesenchymal CTC survival probability | $2 \times 10^{-4}$ | Luzzi et al. (1998) | |
| $\mathcal{P}_{E/M}$ | Partial-EMT CTC survival probability | $6 \times 10^{-4}$ | Luzzi et al. (1998) | |
| $\mathcal{P}_C$ | CTC cluster survival probability | $2.5 \times 10^{-2}$ | Luzzi et al. (1998)<br>Aceto et al. (2014) | |
| $\mathcal{P}_S^1$ | Probability for surviving cells to extravasate to bones | $\sim 0.5461$ | Kuhn Laboratory (2017) | $\sim 0.5461$ |
| $\mathcal{P}_S^2$ | Probability for surviving cells to extravasate to lungs | $\sim 0.2553$ | Kuhn Laboratory (2017) | $\sim 0.2553$ |
| $\mathcal{P}_S^3$ | Probability for surviving cells to extravasate to liver | $\sim 0.1986$ | Kuhn Laboratory (2017) | $\sim 0.1986$ |
| $\mathcal{P}_{EMT}^M$ | Probability for full EMT on $\Omega_P$ | $1 \times 10^{-2}$ | Estimated | |
| $\mathcal{P}_{EMT}^{E/M}$ | Probability for partial EMT on $\Omega_P$ | $2 \times 10^{-2}$ | Estimated | |
| $\mathcal{P}_{EMT}^{E/M*}$ | Additional probability for partial EMT on $\Omega_P$ at tumour edge | 0.15 | Estimated | |
| $\mathcal{P}_{MET}^E$ | Probability for partial MET on $\Omega_S^{1,2,3}$ | 0.5 | Estimated | |
| $w(0, \vec{x}), \vec{x} \in \Omega_P$ | Breast initial ECM density | 1 | ICRP (2009) | $1.020 \text{ g cm}^{-3}$ |
| $w(0, \vec{x}), \vec{x} \in \Omega_S^1$ | Bone initial ECM density | 0.9608 | ICRP (2009) | $0.980 \text{ g cm}^{-3}$ |
| $w(0, \vec{x}), \vec{x} \in \Omega_S^2$ | Lung initial ECM density | 1.0392 | ICRP (2009) | $1.060 \text{ g cm}^{-3}$ |
| $w(0, \vec{x}), \vec{x} \in \Omega_S^3$ | Liver initial ECM density | 1.0294 | ICRP (2009) | $1.050 \text{ g cm}^{-3}$ |

## C. Funding

This work was supported by the Engineering and Physical Sciences Research Council (EPSRC) [to L.C.F.]; EPSRC Grant No. EP/N014642/1 (EPSRC Centre for Multiscale Soft Tissue Mechanics With Application to Heart & Cancer) [to M.A.J.C.].

## Footnotes

1 The notation (*x*_*i*_, *y*_*j*_) ∈ *Ω*_P_ is a result of the discretisation of the grids in our model, as described in detail in Appendix B of Franssen et al. (2019a).

